# Few temporally distributed brain connectivity states predict human cognitive abilities

**DOI:** 10.1101/2022.12.23.521743

**Authors:** Maren H. Wehrheim, Joshua Faskowitz, Olaf Sporns, Christian J. Fiebach, Matthias Kaschube, Kirsten Hilger

## Abstract

Human functional brain connectivity can be temporally decomposed into states of high and low cofluctuation, defined as coactivation of brain regions over time. Rare states of particularly high cofluctuation have been shown to reflect fundamentals of intrinsic functional network architecture and to be highly subject-specific. However, it is unclear whether such network-defining states also contribute to individual variations in cognitive abilities – which strongly rely on the interactions among distributed brain regions. By introducing CMEP, a new eigenvector-based prediction framework, we show that as few as 16 temporally separated time frames (< 1.5% of 10min resting-state fMRI) can significantly predict individual differences in intelligence (*N* = 263, *p* < .001). Against previous expectations, individual’s network-defining time frames of particularly high cofluctuation do not predict intelligence. Multiple functional brain networks contribute to the prediction, and all results replicate in an independent sample (*N* = 831). Our results suggest that although fundamentals of person-specific functional connectomes can be derived from few time frames of highest connectivity, temporally distributed information is necessary to extract information about cognitive abilities. This information is not restricted to specific connectivity states, like network-defining high-cofluctuation states, but rather reflected across the entire length of the brain connectivity time series.

## 1. Introduction

Humans differ in cognitive ability as assessed by measures of intelligence. Individual differences in intelligence, in turn, are associated with important life outcomes like academic achievement (Deary et al., 2007), socio-economic status (Strenze, 2007), or health (e.g., Batty & Deary, 2004). While intact brain functions are a necessary pre-condition for intelligent behavior and thought, as evidenced by neuropsychological data from lesion studies (e.g., Woolgar et al., 2010), the neurobiological mechanisms underlying individual differences in intelligence are not yet fully understood (e.g., Basten et al., 2015; Hilger et al., 2020a). One currently emerging hypothesis is that individual differences in intelligence (or general cognitive ability) are related not only to the structure or function of distinct brain regions, but to their interactions and the information flow between them (e.g., Hilger et al., 2017; Hilger et al., 2020b; Litwińczuk et al., 2022; Ooi et al., 2022; see also Barbey, 2018) – a proposal that is at least partly consistent with psychological theories postulating that general intelligence is the result of the coordination between several fundamental cognitive processes (including, e.g., working memory capacity and mental processing speed; e.g., Duncan et al., 2020; Frischkorn et al., 2019; for review see Euler & McKinney, 2021; Hilger et al., 2022).

In the past few years, abundant functional MRI (fMRI) research has established that the human brain is organized into functionally dissociable large-scale networks (Fox et al., 2005; Greicius et al., 2003; Seeley et al., 2007; Sporns & Betzel, 2016), consisting of anatomically distributed brain regions whose activity covaries across time, even in the absence of specific cognitive task demands (task-free or resting-state fMRI). Multiple studies have demonstrated that individual differences in cognitive ability can be predicted from resting-state functional MRI connectivity (e.g., Cai et al., 2020; Chen et al., 2022; Dubois et al., 2018; Finn et al., 2015; Pervaiz et al., 2020), so that a link between functional brain network organization and individual differences in general cognitive ability (intelligence) is widely accepted (see also Hilger & Sporns, 2021, for an overview of network neuroscience studies on intelligence). The exact nature of this relationship, however, remains to be fully understood.

Human network neuroscience studies have commonly relied on the analysis of ‘static’ intrinsic functional connectivity, which is derived from correlations between the entire BOLD time series of different brain regions (typically assessed with task-free resting-state fMRI measurements of 5 to 10 minutes in length, e.g., Friston et al., 1993). Newer methodological advances made it possible to explore functional connectivity as a ‘dynamic’ property of the human brain (see Lurie et al., 2020; for an overview) and suggest that individual differences in behavior and cognition may be related to variations of functional connectivity over time (e.g., Fong et al., 2019; Hilger et al., 2020b; Jiang et al., 2020). Most recently, time-resolved analyses of intrinsic functional connectivity have demonstrated that a small fraction of the fMRI BOLD time series is characterized by particularly strong functional interactions between brain regions (Cifre et al., 2020; Liu & Duyn, 2013; Tagliazucchi et al., 2012) and sufficient to account for several fundamental properties of the static network architecture, including the overall (time-averaged) configuration of connectivity patterns and their hierarchical modular structure (Esfahlani et al., 2020). These ‘network-defining’ states of high brain-wide cofluctuation (top 5% of the entire timeseries, i.e., 55 out of 1,100 time frames in Esfahlani et al., 2020) were suggested to be temporally stable and highly individual-specific (Betzel et al., 2022; Cutts et al., 2023; Sporns et al., 2021), as subjects can be identified (in the sense of ‘network fingerprinting’; cf. Finn et al., 2015) significantly better on the basis of these high-connectivity states, compared to the same number of low cofluctuation time frames (Esfahlani et al., 2020). Following up on these findings, we here explore whether individual differences in general cognitive ability (intelligence) can be predicted from specific states of intrinsic functional brain connectivity, thereby defining ‘states’ as selections of fMRI time frames characterized by a particular selection criterion (such as strongest vs. weakest brain-wide cofluctuation).

To this end, we investigated the association between general intelligence and different states of intrinsic functional connectivity in a sample of 263 adults for which intelligence test data (Full-Scale Intelligence Quotient, FSIQ, from the Wechsler Abbreviated Scale of Intelligence; WASI; Wechsler, 1999) and temporally highly resolved resting-state fMRI data are available (NKI Enhanced Rockland Sample; Nooner et al., 2012). To increase the robustness of prediction results we developed a cross-validated machine learning-based prediction framework that circumvents the necessity of selecting arbitrary statistical threshold parameters. First, we replicated earlier reports that static (time-averaged) functional connectivity can significantly predict intelligence. We then identified high cofluctuation time frames as described by Esfahlani et al., (2020) and replicate their finding that the general structure of static functional connectivity is strongly determined by a small number of such high cofluctuation states. However, intelligence could not be predicted from these ‘network-defining’ states, nor from an equal number of time frames reflecting particularly low brain-wide cofluctuation. Systematic analyses showed that more independent (i.e., temporally separated) time frames are required to predict intelligence than to recover major characteristics of general functional network structure. However, given temporal independence, this number can be as low as 16 time frames and appears largely independent of the ability to predict global network features. Finally, multiple functional networks were involved in the prediction, suggesting intelligence as whole-brain phenomenon. All results have been replicated in an independent sample (*N* = 812, Van Essen et al., 2013).

## 2. Materials and Methods

### 2.1. Participants

#### 2.1.1. Primary sample

Data from the Enhanced NKI Rockland sample (NKI-RS Enhanced Sample; acquired by the Nathan S. Kline Institute for Psychiatric Research, Release 1-5; Nooner et al., 2012) were used in all analyses. The NKI dataset was selected as primary sample, because a) participants were characterized by a well-established and reliable measure of intelligence (the Full Scale Intelligence Quotient, FSIQ; Wechsler, 1999), which served as estimate of general cognitive ability, b) the sample is characterized by a broad variation in age and intelligence (in contrast to, e.g., to the Human Connectome Project; see 2.1.2. Replication sample), as it was explicitly recruited to be representative of the population. We restricted our analyses to subjects for whom complete fast-sampling fMRI data and IQ scores were available and who passed the Connectome Computational System (CCS) quality check (implying exact motion thresholds; see below), the fmriPrep quality control as well as our visual inspection. This resulted in a final sample of *N* = 263 participants (172 females, 231 right-handed, mean age: 47.14, 18 – 83 years; FSIQ: mean 105.93, range 74 – 143).

#### 2.1.2. Replication sample

The generalizability of our findings to an independent sample was assessed with data from the Human Connectome Project (HCP; 1200 release; Van Essen et al., 2013). After removing subjects with incomplete MRI data, missing phenotypic measures, or more than 10% motion spikes in the fMRI data (defined as framewise displacement, FD > .25 mm; see Parkes et al., 2018), the replication sample comprised *N* = 831 subjects (390 males, 756 right-handed, mean age: 28.55, 22 – 36 years). In contrast to the primary sample, the HCP dataset did not contain an intelligence test, but the Penn Matrix Test (PMAT) as a fast-to-administer approximation of fluid intelligence (Bilker et al., 2012). PMAT scores represent the number of correct responses out of 24 items (mean 17.32, range 5 – 24 in our sample), but were not normally distributed in our sample (Shapiro-Wilk-Test: *W* = .92, *p* < .001). To construct a more comprehensive measure of general cognitive ability, we used 12 cognitive performance scores (Barch et al., 2013) to calculate a latent factor with bifactor analysis (Dubois et al., 2018). Based on one of the most influential theories of intelligence (Spearman, 1904), such a ‘*g*-factor’ constitutes a valid representation of general cognitive ability. The standardized estimate of the *g*-factor (mean 0; range −3 – 2.32) was used as variable of interest for replication analyses.

All study procedures were approved by the NKI Institutional Review Board (#239708; primary sample), the Washington University Institutional Review Board (replication sample, for details see Van Essen et al., 2013), and informed consent in accordance with the declaration of Helsinki was obtained from all participants.

### 2.2. Data acquisition and preprocessing

#### 2.2.1. Primary sample

Fast sampling resting-state fMRI data (9:15 min; eyes open; TR = 645 ms, 860 time frames, voxel size = 3 mm isotropic, 40 slices, TE = 30 ms, flip angle = 60°, FOV = 222 x 222 mm^2^) was obtained with a 32-channel head coil on a 3T Siemens Tim Trio scanner. A structural scan (T1-weighted, voxel size = 1 mm isotropic, 176 slices, TR = 1,900 ms, TE = 2.52 ms, flip angle = 9°, FOV = 250 × 250 mm^2^) was acquired for coregistration. For subjects with more than one T1-weighted (T1w) image, an unbiased subject-specific average was constructed using ANT’s antsMultivariateTemplate Construction2.sh script. All T1w were denoised using ANT’s DenoiseImage function. We then preprocessed the fMRI data in an initial step with fMRIPrep, followed by second step to remove nuisance confounds from the data with Nilearn.

For the first fMRI preprocessing step we used fMRIPrep version 20.2.5 (Esteban et al., 2019), a Nipype (Gorgolewski et al., 2011) based tool, involving the following steps (description based on freely distributed text describing analysis steps performed by fMRIPrep): Each T1w (T1-weighted) volume was corrected for intensity non-uniformity using N4BiasFieldCorrection v2.1.0 (Tustison et al., 2010) and skull-stripped using antsBrainExtraction.sh v2.1.0 (using the NKI template). Brain surfaces were reconstructed using recon-all from FreeSurfer v6.0.1 (Dale et al., 1999), and the brain mask estimated previously was refined with a custom variation of the method to reconcile ANTs-derived and FreeSurfer-derived segmentations of the cortical gray-matter of Mindboggle (Klein et al., 2017). Spatial normalization to the ICBM 152 Nonlinear Asymmetrical template version 2009c (Fonov et al., 2009) was performed through nonlinear registration with the antsRegistration tool (ANTs v2.1.0; Avants et al., 2008), using brain-extracted versions of both the T1w volume and the template. Brain tissue segmentation of cerebrospinal fluid (CSF), white-matter (WM) and gray-matter (GM) was performed on the brain-extracted T1w using FAST (FSL v5.0.9; Zhang et al., 2001). Functional data was slice time corrected using 3dTshift from AFNI v16.2.07 (Cox, 2012) and motion corrected using MCFLIRT (FSL v5.0.9; Jenkinson et al., 2002). “Fieldmap-less” distortion correction was performed by co-registering the functional image to the same-subject’s T1w image with intensity inverted (Huntenburg et al., 2014; Wang et al., 2017) constrained with an average fieldmap template (Treiber et al., 2016), implemented with antsRegistration (ANTs). This was followed by co-registration to the corresponding T1w using boundary-based registration (Greve & Fischl, 2009) with nine degrees of freedom, using bbregister (FreeSurfer v6.0.1). Motion correcting transformations, field distortion correcting warp, BOLD-to-T1w transformation and T1w-to-template (MNI) warp were concatenated and applied in a single step using antsApplyTransforms (ANTs v2.1.0) with Lanczos interpolation.

Two physiological noise regressors were extracted by estimating principal components with CompCor (Behzadi et al., 2007), i.e., temporal (tCompCor) and anatomical (aCompCor). A mask to exclude signal with cortical origin was then obtained by eroding the brain mask, ensuring that it only contains subcortical structures. Afterwards, six tCompCor components were calculated including only the top 5% temporally variable voxels within that subcortical mask. For aCompCor, six components were constructed within the intersection of the subcortical mask and the union of CSF and WM masks calculated in T1w space, after their projection to the native space of each functional run. Frame-wise displacement (Power et al., 2014) was computed for each functional run using the implementation of Nipype. For more details of the pipeline see https://fmriprep.readthedocs.io/en/20.2.5/workflows.html.

After the initial preprocessing with fMRIPrep, subject-specific functional connectivity matrices were constructed. Therefore, brain volumes were parcellated into 200 regions of the Schaefer atlas (Schaefer et al., 2018), which allow for the assignment of brain regions to seven functional networks: VIS, visual network; SMN, somatomotor network; DAN, dorsal attention network; VAN, ventral attention network; LIM, limbic network; CON, control network; DMN, default mode network. Note that the Schaefer 200 parcellation was rendered in each subject’s volumetric anatomical space. In detail, FreeSurfer’s surface warp was applied to bring the parcellation from fsaverage space to the subject space, using the mris_ca_label tool, in conjunction with a pre-trained Gaussian classifier surface atlas. This process ensures that the parcellation is rendered in the grey matter ribbon, and that the parcellation was non-linearly warped to each subject based on individual curvature and surface patterns. Each functional image in anatomical space was linearly detrended, band-pass filtered (0.008–0.08 Hz), nuisance regressed, and standardized using Nilearn’s signal.clean function, which removes confounds orthogonally to the temporal filters. The nuisance regression strategy included six motion estimates, mean signal from a white matter, cerebrospinal fluid, and whole brain mask, derivatives of these nine regressors, and squares of the respective 18 terms. Following these preprocessing operations, the mean signal was then taken at each node within the subject’s volumetrically rendered parcellation. The resultant time series were then z-scored across time and the first and last 20 frames were removed.

For each participant separately, functional connections were then defined as weighted undirected edges and modelled using Fisher *z*-transformed Pearson correlation coefficients between these z-scored time series of each pair of brain regions (Fig. 1a, d). These edges represent brain connectivity as averaged across the whole duration of the resting-state scan and provide the basis for the construction of each participant’s static connectivity matrix. In addition to the motion correction during preprocessing, we controlled for potential remaining influences of head motion by regressing out mean FD from the FSIQ scores as well as from each connectivity value with linear regression (http://scikitlearn.org/stable/modules/generated/sklearn.linear_model.LinearRegression.html).

**Fig 1.**
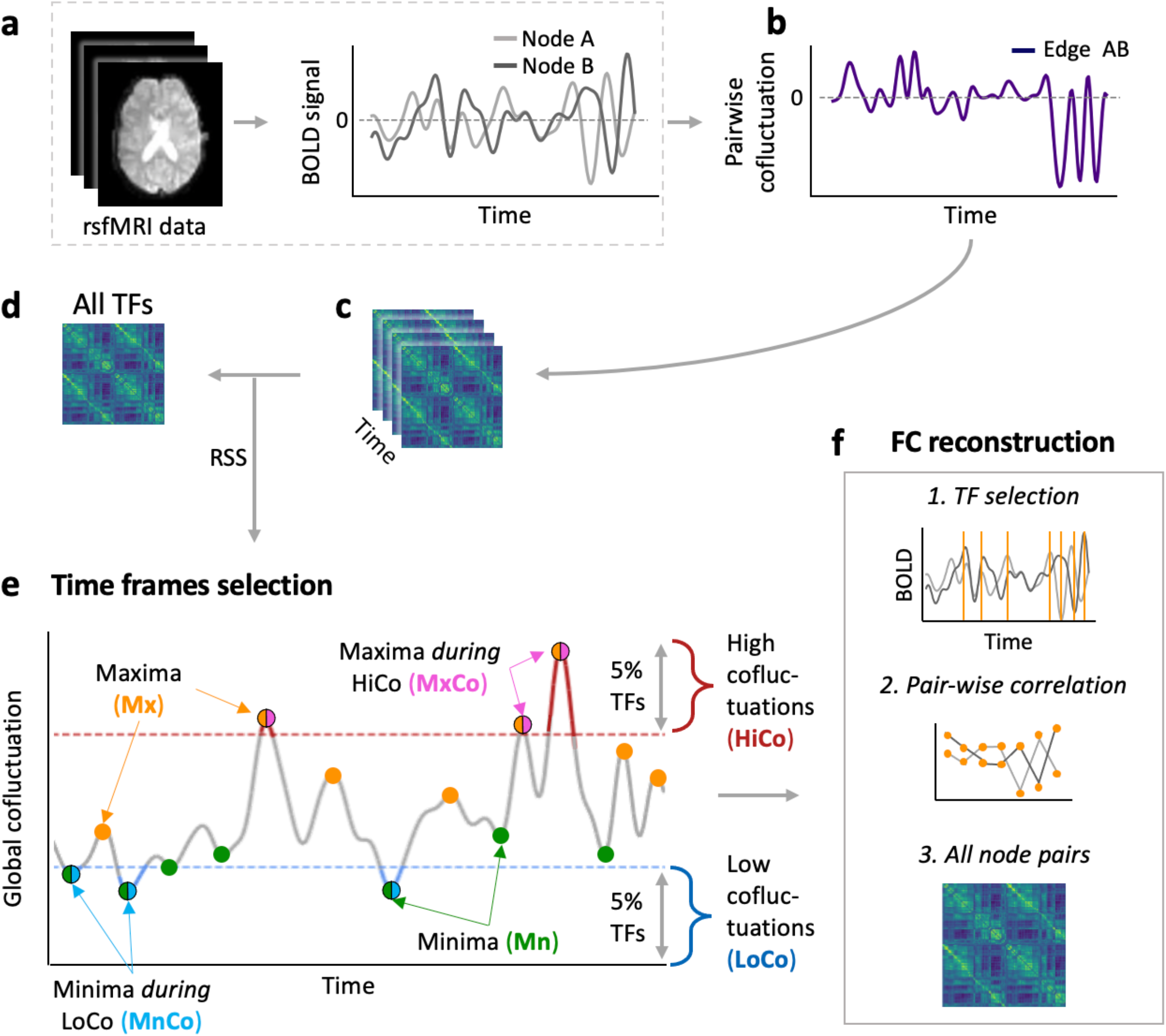
Schematic illustration of analysis steps to derive different types of state-restricted functional connectivity. (**a**) Resting-state fMRI data were parcellated into 200 functional brain regions (nodes; Schaefer et al., 2018) and node-specific BOLD activation time courses were extracted. (**b**) For a given node pair, the strength of their cofluctuation is given by the product of their *z*-scored neural activation time series. Time-resolved whole-brain connectivity matrices (**c**) were then computed on the basis of all node-pairs instantaneous cofluctuation. Averaging this time-resolved connectivity matrices across all time frames (TFs) yields the full (‘static’) functional connectivity matrix (**d**). (**e**) The strength of instantaneous brain-wide (global) cofluctuation is computed as the root-sum-square (RSS) of all node-pair cofluctuations (Esfahlani et al., 2020). Based on this global cofluctuation time series, six different brain connectivity states were defined (see Methods for details). The selections comprise i) the 43 time frames with the highest/lowest values of global cofluctuation (HiCo/LoCo), ii) only the maxima/minima within each of these short episodes of highest/lowest cofluctuation states (MxCo/MnCo, < 16 TFs) and iii) a larger set of, again, 43 time frames, but separated in time and representing the highest maxima/lowest minima within the complete RSS time series (maxima/minima, Mx/Mn). Apart from these specific selections of time frames, we also varied the number of frames and studied selections based on randomly drawn time-points. (**f**) On the basis of these different selections, functional connectivity matrices were computed as Fisher-z transformed Pearson correlation representing the six subject-specific brain connectivity states. rsfMRI, resting-state functional magnetic resonance imaging.

#### 2.2.2. Replication sample

Data of the replication sample consisted of the four resting-state runs from the HCP (15 min, 1,200 time points each; for details of data acquisition see Van Essen et al., 2013). The scanning parameters were: voxel size = 2 mm isotropic, 72 slices, TR = 720 ms, TE = 33 ms, flip angle = 52°, FOV = 208 x 180 mm^2^ (for details Smith et al., 2013). We obtained the minimally preprocessed data from the HCP (Glasser et al., 2013), involving an initial correction for head motion and B0 distortion, co-registration to T1-weighted structural images, and normalization to MNI152 space (i.e., the MNI template included in the FSL package and published as part of the HCP pipelines: https://github.com/Washington-University/HCPpipelines/tree/ master/global/templates). For further preprocessing, we followed strategy number six (24HMP+8Phys+4GSR) described in Parkes et al. (2018) which has been shown to perform well on the HCP resting-state data and maintains the same temporal degrees of freedom for all subjects. This strategy comprises regressing out a) 24 motion parameters including the raw scores as well as the squares of both the original and the derivate time series (Satterthwaite et al., 2013), b) eight physical parameters including white matter (WM) and cerebrospinal fluid (CSF) signals along with their temporal derivatives, squares, and squares of derivatives, and c) the global signal with its temporal derivative, square term, and the square of the derivatives. Also, a temporal band-pass filter of .008-.08 Hz was applied (Parkes et al., 2018). Note that for comparability with the primary sample, i.e., to ensure same number of time frames, only 860 time frames in the center of each of the four resting-state time series were used for further analyses. Brain volumes were parcellated into 100 regions of the Schaefer atlas (Schaefer et al., 2018), which allow direct alignment to the seven Yeo networks also used in the primary sample (Yeo et al., 2011). Again, for each individual static (time-averaged) connectivity matrix, weighted edges were modelled on the basis of Fisher *z*-transformed Pearson correlation coefficients and remaining influences of head motion were controlled via regressing out mean FD and the number of spikes. All further analyses were computed for each resting-state run separately, i.e., four times.

### 2.3. Time-resolved brain connectivity analyses

Following Esfahlani et al. (2020), we resolved connectivity in time to capture the strength of brain-wide cofluctuation for each time frame of the BOLD time series. Consider the Pearson correlation coefficient *r_ij_* that reflects the time-averaged strength of interaction (functional connectivity) between two given nodes *i* and *j:*

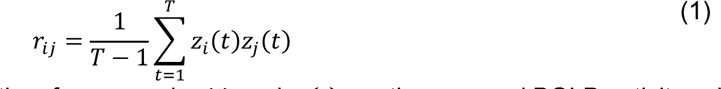

where *T* is the number of fMRI time frames and *z*_i_(*t*) and *z*_j_(*t*) are the z-scored BOLD activity values at time *t* of nodes *i* and *j*, respectively. For example, for node *i*, the z-scored BOLD activity is defined as

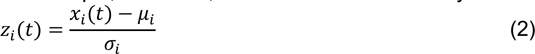

where *x*_i_(*t*) represents the BOLD activity at time *t*, μ_i_ is the mean of node *i*’s time series, and σ_i_ is the respective standard deviation. The instantaneous cofluctuation between node *i* and node *j* at time frame *t* (representing time-resolved functional connectivity; Fig. 1b) is thus given by:

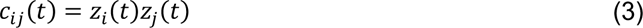

On the basis of *c*_ij_, we computed a three-dimensional time-resolved connectivity matrix for each participant (see Fig. 1c for a schematic illustration) with the shape *n* x *n* x *T*, where *n* is the number of nodes in the respective fMRI dataset (NKI: *n* = 200, HCP: *n* = 100) and *T* is the number of time frames (*T* = 860). The strength of brain-wide cofluctuation per time frame was then quantified by the root-sum-square (RSS; Fig 1e) of the cofluctuation values of all node pairs *c*_ij_(*t*) at a given time frame *t*:

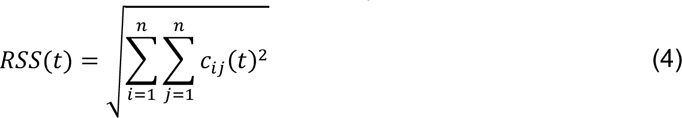

On the basis of this RSS, three different types of connectomes (each relying on different selections of time frames) were created for each participant: A) In line with Esfahlani et al. (2020), we defined high cofluctuation states (also referred to as ‘events’) as those time frames with the 5% highest RSS values and low cofluctuation states as time frames with the 5% lowest RSS values, respectively (in our samples 43 time frames; Fig 1e, HiCo/LoCo, red and blue parts of the RSS curve). B) As the temporally adjacent time frames included in these high and low cofluctuation states events are highly correlated with each other (for discussion see Ladwig et al., 2022), we replaced each event of temporally grouped highest/lowest cofluctuation frames by the single frame with maximal/minimal cofluctuation within this event, thereby reducing the total number of frames considerably (Fig 1e, MxCo/MnCo, pink and light blue dots). C) Next, we selected from all maxima/minima within the entire RSS time series the 43 time frames with the overall highest/lowest cofluctuation values (corresponding to again 5% of data; Fig 1e, Mx/Mn, orange and green dots). Note that case A and C include the same number of time frames, whereas case B represents the intersection between the frames of case A and C. In case B and C, but not in case A, the selected time frames were temporally separated from each other. For further analyses, and additionally to the previously computed static functional connectivity based on all time points (see above; Fig 1d), functional connectivity matrices (Fisher z-transformed Pearson correlation, Fig. 1f) were computed from each of these six selected subsets of time frames (HiCo/LoCo, MxCo/MnCo, Mx/Mn), as well as for comparable random uniformly drawn subsets of time points. The resulting individual-specific functional connectivity matrices (seven per subject) were used as input to prediction analyses.

### 2.4. Covariance maximizing eigenvector-based predictive modelling (CMEP)

We developed a two-staged machine learning-based predictive modelling framework, which we refer to as covariance maximizing eigenvector-based predictive modelling (CMEP, see Fig. 2 for a schematic illustration). The basic idea of CMEP is to first create candidate features (based on eigenvectors) within a training set that share a strong linear relationship (covariance) with the target variable of interest, and then, in a second step, to assess the predictive power of these features by training and testing a prediction model (ElasticNet regression) based on these features. Notably, the ElasticNet regression circumvents the need to select features based on arbitrary threshold parameters (as is the case in many other prediction frameworks used for brain imaging data).

**Fig. 2.**
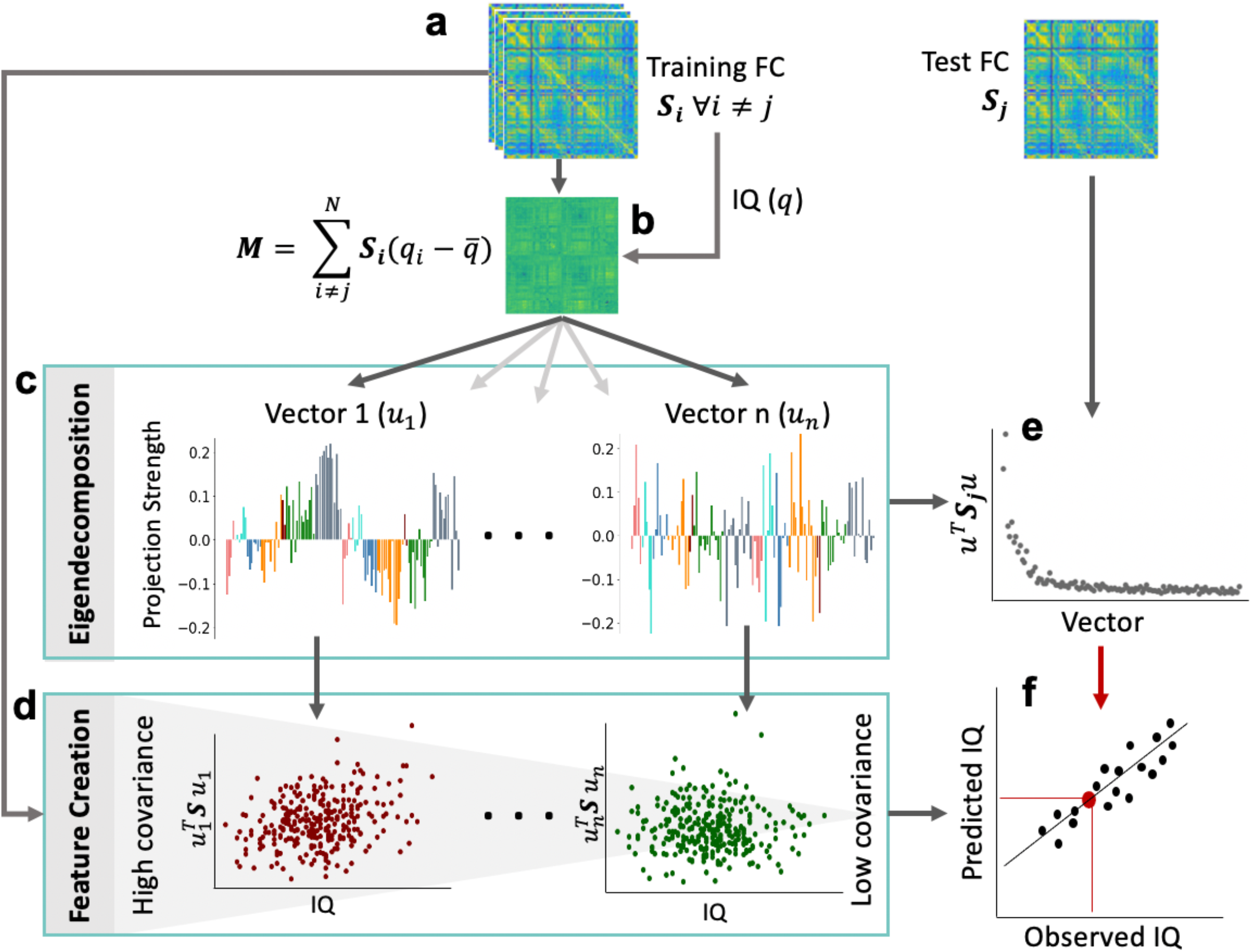
Schematic illustration of the Covariance Maximizing Eigenvector-Based Predictive Modelling (CMEP) framework. (**a**) For leave-one-out cross validation the functional connectivity ***(S)*** was computed for every subject *i* and these data were split into a training and a test set. (**b**) Calculation of an IQ-weighted group connectivity matrix ***(M)*** to enhance differences in intelligence-related features in the training sample. (**c**) Eigendecomposition of ***M*** generates the eigenvectors (*u*_1_ … *u*_n_). The entries of the eigenvectors were highlighted in different colors according to the seven functional brain networks they correspond to (Yeo et al., 2011; within each hemisphere; networks from left to right: visual, somatomotor, dorsal attention, ventral attention, limbic, control, and default mode). Projecting subject-specific functional connectivity onto these eigenvectors generates brain connectivity features for the training (**d**) and the left-out test sample (**e**). (**f**) All training set features were used to fit an ElasticNet regression and tested for generalization with the withheld test set features. Model performance was assessed by comparing the predicted with the observed intelligence scores. See Methods for further details.

#### 2.4.1. Feature construction

The key characteristic that differentiates CMEP from previous predictive modelling approaches is its feature construction method which amplifies features whose expression covary with a target variable (here intelligence). To this aim, functional connectivity matrices are projected into a scalar space chosen such that the covariation between individual intelligence scores and this scalar projection is maximal.

More specifically, within the training set, we find a projection 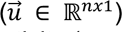 that maximizes the covariance between intelligence scores 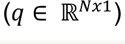 and functional connectivity 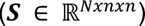:

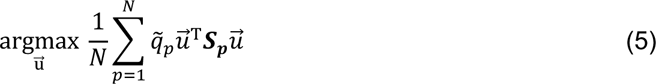

where 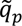 represents the mean adjusted intelligence score, i.e., individual IQ (*q_p_*.) minus group-average 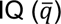 of a participant *p*. As the projection 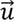 is independent of *p*, Eq. 5 can be reformulated as follows

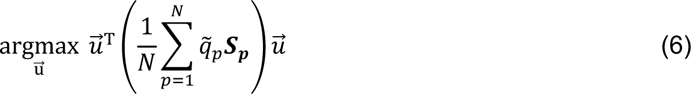

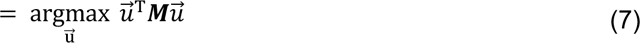

The matrix ***M*** (Fig. 2a,b) emphasizes intelligence-related differences in functional connectivity within the training set. Note that the leading eigenvector of ***M*** corresponds to the feature that maximally covaries with intelligence. However, it turned out that the prediction of general intelligence was more robust when including also other eigenvectors. Therefore, we next calculated the eigendecomposition of the general intelligence-weighted matrix ***M***:

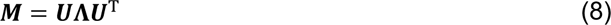

where ***U*** is the matrix of eigenvectors 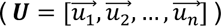 and Λ is the diagonal matrix of the corresponding eigenvalues in decreasing order (Fig. 2c). We assumed that ***M*** is full rank and since it is real and symmetric, all eigenvalues are also real and the eigenvectors orthogonal. Note that the entries within a given eigenvector are interpretable as the weights of the corresponding nodes for that particular vector (Fig. 2c).

Then, subject-specific features are derived by projecting each participant’s connectivity matrix ***S***_p_ onto the previously computed eigenvectors 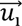:

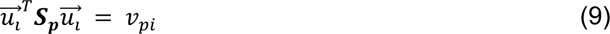

This step results in an individual feature vector 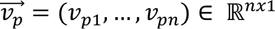 for each participant *p* (Fig. 2d and f). The number of features is defined by the number of eigenvectors 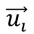 and is thus equal to the number of nodes (*n*), i.e., 200 in the primary sample and 100 in the replication sample. Finally, an ElasticNet regression model is trained and optimized to predict intelligence scores from the generated feature vectors of an independent test set (Fig. 2e,f).

#### 2.4.2. Prediction framework

The feature construction step of CMEP is embedded within a cross-validated prediction framework in which, within each cross-validation fold, a new set of eigenvector-based features are created to ensure strict independence between training and test set. Specifically, the prediction framework consists of two nested cross-validation loops. In the outer loop (primary sample: leave-one-subject-out cross validation; replication sample: leave-one-family-out due to family relations between participants) first, the eigenvectors ***U*** are computed from the training set. Second, the feature vectors 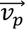 are constructed for each individual subject and third, an ElasticNet regression model is trained to predict the intelligence scores (*q_p_*) from the subject-specific feature vectors 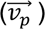 within the training set. The hyperparameters of the ElasticNet model are optimized within the *inner* loop (5-fold cross-validation of the training set). ElasticNet regularizes a linear regression model via the L1 (Lasso; favoring feature sparsity) and L2 (Ridge; encouraging coefficient shrinkage) norm to avoid overfitting (Zou & Hastie, 2005). Therefore, ElasticNet regression allows to utilize all created features as candidates for the prediction without manually setting an a-priori selection threshold. The model is formalized as:

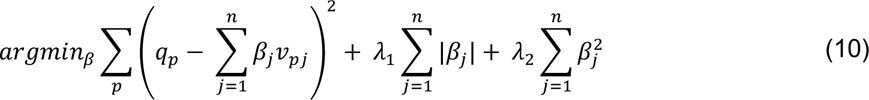

where β is the vector of regression weights and *q_p_*. and *v_p_*. are the cognitive ability score and the feature vector, respectively, of this specific participant. The hyperparameters λ_1_ (L1-penalty) and λ_2’_ (L2-penalty) are optimized to minimize the mean squared error (MSE) between observed and predicted intelligence scores. The sklearn ElasticNetCV implementation in Python was used with parameter choices for 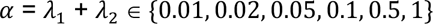, and 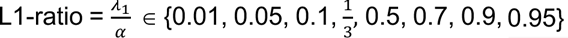 to reduce computational costs. The obtained regression weights are then applied to the feature vector of the held-out subject (test set) to predict their intelligence score.

Prediction performance in terms of difference between predicted and observed intelligence scores of the testing samples (in leave-one-subject-out cross validation: average across the predictions after leaving each subject out once) was evaluated using Pearson correlation coefficient (*r*), mean squared error (MSE), root mean squared error (RMSE), and mean absolute error (MAE). While MSE and RMSE capture differences in bias and precision, correlation coefficients and MAE can be more meaningfully interpreted and allow for direct comparability with previous reports (Hilger, et al., 2020a).

#### 2.4.3. Prediction robustness analyses

Three different control analyses were conducted with static (time-averaged) connectivity matrices to assess the validity of CMEP. Specifically, the robustness of prediction derived from CMEP were compared with those derived from the most frequently used neuroscientific prediction framework, i.e., connectome-based predictive modeling (CPM; Finn et al., 2015). First, we assessed the robustness across different data set splits by randomly (100 times) dividing the data into 10 cross validation splits. Second, to test the robustness across different samples sizes, the training sample was randomly (100 times) reduced to 10% of the original data size within each iteration of a stratified 10-fold cross-validation. Finally, the transferability of each model was assessed by training it on one sample (e.g., the primary sample, NKI) and testing it on an independent sample (e.g., the replication sample, HCP). To this end, both samples were parcellated into the 200 nodes partition and all intelligence scores were standardized. 100 different models were fitted, each with a randomly bootstrapped composition of the training set. All analyses described above were conducted four times, i.e., based on CMEP, the positive CPM network, the negative CPM network, and the combination of both. Specifically, in CPM prediction features are calculated as the sum over all functional connections that are significantly correlated with intelligence above a given threshold (here: *p* < .001; for details see Finn et al., 2015; Shen et al., 2017). Positively correlated connections were used to predict from a positive network, negatively correlated connections to predict from a negative network, and the positive and negative networks to predict from the whole brain.

#### 2.4.4. Significance tests

Non-parametric permutation tests were used to assess the statistical significance of above-chance predictive model performance. Specifically, we took all *N* (primary sample: 263; replication sample: 831) target values (cognitive ability scores) and permuted them, which resulted in a random assignment between subjects and target values, and then assessed prediction performance (*r*, MSE, RMSE, MAE). This step was repeated 1,000 times. The significance of a prediction model (*p*-value) was then calculated as fraction of times the model’s performance of predicting the permuted scores was better than the performance of predicting the actually observed scores. In cases where the prediction performance of two models was compared, a similar non-parametric procedure was adopted by comparing the difference in prediction performance between both models trained with the true targets and the difference in performance with permuted targets. Statistical significance was accepted for *p* values < .05.

### 2.5. Data Availability Statement

Data of the primary sample (NKI) can be accessed under http://fcon_1000.Projects.nitrc.org/indi/enhanced/. Data of the replication sample (HCP) can be downloaded from https://www.humanconnectome.org/study/hcp-young-adult. The analyses code for the preprocessing of the NKI sample can be obtained from https://fmriprep.readthedocs.io/en/20.2.5/workflows.html and the analyses code for further preprocessing of the minimally preprocessed HCP data (replication sample) is available under https://github.com/faskowit/app-fmri-2-mat. Code of all further analyses (including CMEP) has been deposited on GitHub (https://github.com/kaschube-lab/CMEP).

## 3. Results

General cognitive ability was assessed with an established measure of intelligence (FSIQ from the WASI; Wechsler, 1999), and descriptive statistics show that the intelligence quotient is normally distributed in our sample of 18 to 83 year old adults (172 females; Fig. S1). Time-resolved functional connectivity between brain regions (network nodes) was extracted from fast-sampling resting-state fMRI data (primary sample: 200 regions, TR = 645 ms; replication sample: 100 regions, TR = 720 ms; see Methods and Fig. 1a,b). A covariance maximizing eigenvector-based predictive modelling framework (CMEP, see Fig. 2 and Methods for details) was developed and used for all prediction analyses. This machine learning-based prediction framework involves eigenvector-based generation of features from functional brain connections that highly covary with the variable of interest (Fig. 2a-e), as well as ElasticNet regression and leave-one-out (LOO) cross validation (Fig. 2f).

### 3.1. Static functional connectivity predicts intelligence

We first used the CMEP prediction framework to replicate previous findings (e.g., Dubois et al., 2018; Finn et al., 2015) that time-averaged (‘static’) functional connectivity derived from the full fMRI time series (Fig. 1d) significantly predicts intelligence (correlation between predicted and observed intelligence scores: *r* = .34; prediction error: MSE = 150.68; all *p* < .001; see Tab. 1).

**Table 1:** Prediction of intelligence from functional brain connectivity

| | TFs | Reconstruction similarity | $r$ | MSE | RMSE | MAE |
| --- | --- | --- | --- | --- | --- | --- |
| Static functional connectivity | 860 | n/a | .34** | 150.68** | 12.28** | 9.75** |
| Highest cofluctuations (HiCo) | 43 | .80 | .02 | 180.76 | 13.45 | 10.83 |
| Lowest cofluctuations (LoCo) | 43 | .54 | .17 | 165.62 | 12.87 | 10.18 |
| Maxima during HiCo (MxCo) | 4-11 | .79 | .17 | 167.70 | 12.95 | 10.30 |
| Minima during LoCo (MnCo) | 4-16 | .40 | .15 | 166.96 | 12.92 | 10.19 |
| Highest maxima (Mx) | 43 | .97 | .43** | 141.00** | 11.87** | 9.34** |
| Lowest minima (Mn) | 43 | .75 | .32** | 153.70** | 12.40** | 9.72** |

*Note:* Covariance maximizing eigenvector-based predictive modeling (CMEP; see Methods) was used in combination with a nested cross-validation scheme (see Methods, Fig. 1 and Fig. 2) to predict individual intelligence scores (Full Scale Intelligence Quotient, FSIQ; WASI, Wechsler, 1999) from static connectivity (all fMRI time frames; TFs), highest and lowest cofluctuations (HiCo/LoCo; 43 TFs), maxima/minima during highest/lowest cofluctuation (MxCo/MnCo; < 16 TFs), and the 43 highest maxima and lowest minima across the whole RSS time series (Mx, Mn; see Methods and Fig. 1). Reconstruction similarity values represent Pearson correlations between the static connectivity matrix (row 1) and the reconstructed connectivity matrix from the respective selection of time frames. Model performance metrics reflect the error between predicted and observed intelligence scores: Pearson correlation coefficient ( $r$ ), mean squared error (MSE), root mean squared error (RMSE), and mean absolute error (MAE). Significance was determined by a non-parametric permutation test with 1,000 iterations and indicated as \*\* if $p < .001$ after Bonferroni correction for 7 comparisons.

To establish the validity of CMEP, we compared its prediction results with those derived from the most frequently used prediction frameworks (connectome-based predictive modelling, CPM; cf, Finn et al., 2015; Shen et al., 2017). Prediction performance of CMEP was superior across different cross-validation splits (Fig. 3a) and when substantially reducing the sample size of the training set to only 10% (Fig. 3b), while most models produced similar results when trained on one sample and tested on a new sample (Fig. 3c). Note that even though CMEP generally performs better for model transferability, all four tested models generate mostly non-significant results. See Fig. S2 for similar robustness results in the replication sample.

**Fig. 3.**
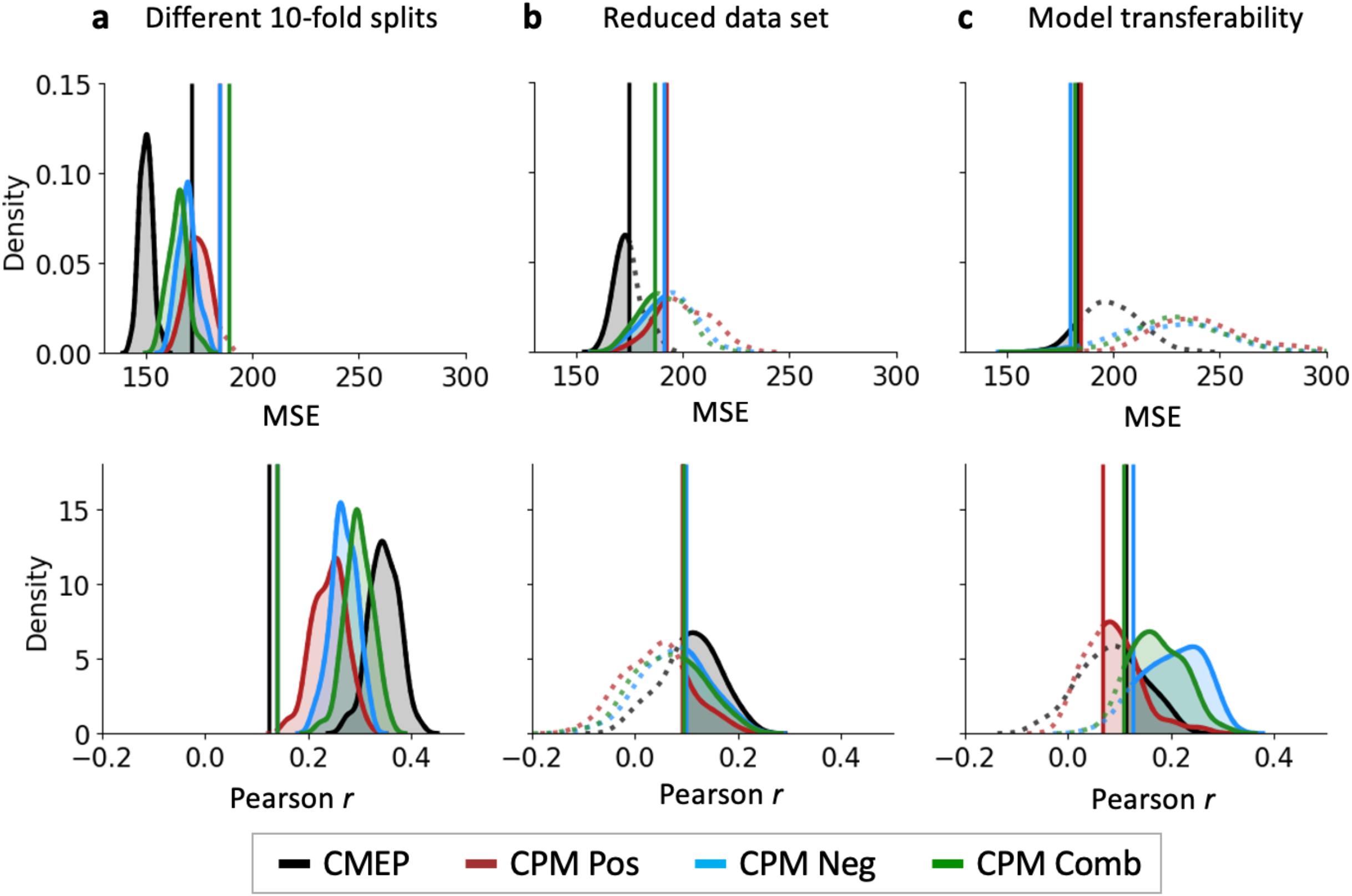
Superior prediction robustness of our method CMEP relative to connectome-based predictive modelling (CPM). Prediction performances (mean squared error, MSE and Pearson correlation between observed and predicted intelligence scores) of CMEP and CPM were computed based on static connectivity (using the entire time series) and three validity analyses (Finn et al., 2015; Shen et al., 2017). All analyses were conducted for CMEP (black, all brain connections), and three CPM prediction pipelines based on positive connections (red), negative connections (blue), and a combination of both (green, all connections). (**a**) Robustness across different data set splits. Data were randomly (100 times) split into 10 folds for cross validation. (**b**) Robustness across different sample sizes. Within stratified 10-fold cross-validation, the training sample was randomly (100 times) reduced to 10% of the original sample size. (**c**) Transferability of the models to a new data set. Models were trained on the replication sample (HCP) and tested on the primary sample (NKI). Both samples were parcellated into the 200 nodes schemata (Yeo et al., 2011) and all intelligence scores were standardized before prediction (but shown on original scale here for better comparability). The training data were randomly bootstrapped (100 times) to account for different compositions of the training data set. The vertical solid lines indicate the significance threshold (p < .05) for each model. Models that were found to be significant are indicated by a shaded area and solid line, insignificant models are depicted with dotted lines.

### 3.2. Network-defining states of highest and lowest cofluctuation are not sufficient to predict intelligence

To test whether intelligence could be predicted from ‘network-defining’ states of highest cofluctuation, we followed the approach introduced by Esfahlani et al. (2020) of analyzing only 5% of the dynamic (time-resolved) functional connectivity time series. First, we operationalized the instantaneous strength of brain-wide (global) connectivity as the root-sum-square (RSS) over the cofluctuation between all pairs of nodes (brain regions), independently for each time frame (see Fig. 1b-e). We then selected the 43 time frames corresponding to the 5% highest global cofluctuation frames (HiCo; Fig. 1e, red), as well as in an independent analysis the 5% time frames with lowest global cofluctuation (LoCo; Fig. 1e, blue). Pearson correlation between the static connectivity matrix (average across all time frames) and the connectivity matrices constructed from only these selections of time frames (Fig. 1f) demonstrated high reconstruction similarity, with on average *r* = .80 (*p* < .001) and *r* = .54 (*p* < .001), respectively, for high and low cofluctuation states (Tab. 1, Fig. 4a). This replicates the previous finding that an individual’s static functional connectome can be well approximated based on just a few time frames of highest cofluctuation (Esfahlani et al., 2020), thus providing an important precondition for all subsequent analyses.

**Fig. 4.**
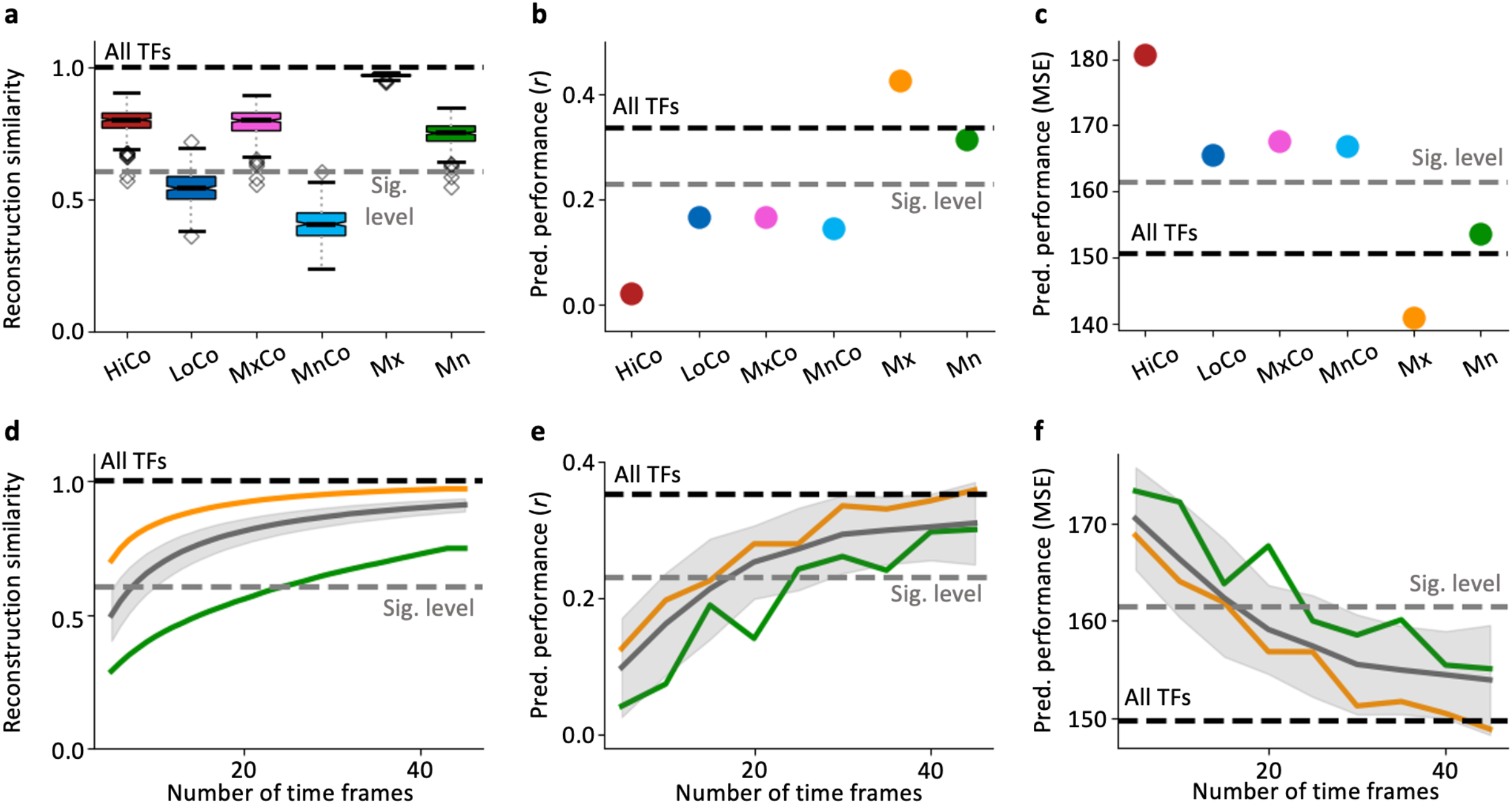
The performance to predict intelligence depends on the number of temporally separated time frames rather than on functional connectivity reconstruction similarity. (**a**) Reconstruction similarity of six different connectivity states operationalized as Pearson correlation between static functional connectivity (constructed from all time frames; TFs) and connectivity matrices reconstructed from six different selections of TFs (see Fig. 1). Boxplots depict the mean and quartiles of the subject-specific reconstruction similarity for all different connectivity states. The whiskers show the 1.5 x interquartile ranges. Outliers are represented by diamonds. (**b, c**) Performance to predict intelligence (FSIQ; WASI, Wechsler, 1999) for the six different connectivity states from using the CMEP prediction framework (see Fig. 2), operationalized in (**b**) as Pearson correlation between predicted and observed scores (*r*) and in (**c**) as mean squared error (MSE). Reconstruction similarity (**d**) and performance to predict intelligence (correlation, **e**; MSE, **f**) as function of the number of time frames comprising cofluctuation maxima or cofluctuation minima (orange or green dots in Fig. 1e). Gray lines represent reconstruction similarity (**d**) and prediction performance (**e**, **f**) from randomly selected time frames. Note that for prediction performances only the two cases are illustrated that allow for significant prediction of intelligence, i.e., 43 highest maxima, Mx; 43 lowest minima, Mn. The upper bounds (black dashed lines) represent reconstruction similarity (**a, d**) or prediction performance (**b, c, e, f**) using all TFs. The lower grey dashed line reflects the approximate 5% significance level (determined as average over all seven models’ significance levels) of the within-subject similarity of static functional connectivity (**a, d**) or intelligence prediction performance (**b, c, e, f**; see Methods). HiCo, highest cofluctuations; LoCo, lowest cofluctuations; MxCo, maxima during HiCo; MnCo, minima during LoCo; Mx, 43 highest maxima; Mn, 43 lowest minima (see also Fig. 1e).

Next, we tested whether these states of highest cofluctuation are predictive of individual differences in intelligence. However, this was not the case (*r* = .02, MSE = 180.76, all *p* > .05). Further, the prediction performance was not significantly different from states of lowest cofluctuation (*r* = .17, MSE = 165.62, all *p* > .05; see Tab. 1; Fig. 4b, red and blue). Comparably low prediction performance was also evident when using an alternative prediction model (CPM; Fig. S3). These results suggest that reconstruction similarity (see previous paragraph) is not a faithful indicator for the ability to extract information about individual differences in cognitive ability from brain connectivity states, and that the ability to predict intelligence depends on a richer set of features than that required for approximating global properties of network structure (Fig. 4a,b).

Notably, these network-defining states of highest and lowest cofluctuation contain a comparably large number of temporally adjacent and thus autocorrelated time frames and comprise only few spatially distinctive coactivation patterns (CAPs; Liu et al., 2018; see Fig. S4). To test whether the low prediction performance of highest and lowest cofluctuation states is due to the small number of temporally independent data points contained in these selections, we next selected only the individual-specific maxima/minima from the previously examined states of highest/lowest cofluctuation, respectively (Fig. 1e; maxima: MxCo pink; minima: MnCo light blue). This strongly reduces the amount of time frames (MxCo; 4-11 TFs, mean: 7.54; MnCo; 4-16 TFs, mean: 9.54), but ensures their temporal separation. Notably, neither reconstruction similarity nor prediction performance decreased markedly (MxCo: reconstruction similarity *r* = .79, *p* < .001; intelligence prediction *r* = .17, MSE = 167.70, all *p* > .05; MnCo: reconstruction similarity *r* = .40, *p* < 001; intelligence prediction *r* = .15, MSE = 166.96, all *p* > .05; Tab. 1; Fig. 4a,b). The same results were observed when using alternative prediction models, but CMEP provides the highest robustness of results (Fig. S3). Together, these analyses suggest that the low predictive power of connectivity states of highest/lowest cofluctuation for intelligence is most likely due to the small amount of independent information included in these selections of temporally adjacent time frames.

### 3.3. A small selection of temporally separated time frames predicts intelligence

The selection of only the maxima/minima during high/low cofluctuation (MxCo/MnCo, see above) indicated that discarding more than half of the time frames from the selection of HiCo/LoCo time frames did neither significantly affect the reconstruction similarity nor the performance to predict intelligence – an observation that points to critical relevance of temporal independence between the selected frames. In a next analysis we therefore selected the 43 highest maxima (Mx) as well as the 43 lowest minima (Mn) across the entire cofluctuation time series (summing up to again 5% of data). This allows for a fairer comparison between the temporally correlated HiCo/LoCo states and the same number of temporally independent states (Mx, Mn). The Mx and Mn states were characterized by a significantly higher number of individual coactivation patterns than the selection of autocorrelated highest/lowest cofluctuation states (HiCo/LoCo), reflecting more independent information within the Mx and Mn states (see Fig. S4). The prediction performance for intelligence increased markedly compared to the previous analyses. Specifically, the 43 highest maxima allowed to reconstruct static functional connectivity on average with *r* = .97 (*p* < .001; Fig. 4a orange) and to significantly predict individual intelligence scores (*r* = .43, MSE = 141.00, all *p* < .001; Tab. 1, Fig. 4b,c orange dot). The same analyses based on a selection of the 43 lowest minima resulted in a reconstruction similarity of *r* = .75 (*p* < .001; Fig. 4a green) and also in significant prediction of intelligence, albeit with a lower prediction performance (*r* = .32, MSE = 153.70, all *p* < .001; Tab. 1, Fig. 4b,c green dot). Again, these results were similar for alternative prediction models, with CMEP showing robustly the best prediction results (Fig. S3).

### 3.4. How much time frames are required to predict intelligence?

To reveal the minimal amount of data necessary to predict individual differences in intelligence, we systematically varied the number of highest maxima and lowest minima time frames, respectively, and investigated how reconstruction similarity and prediction performances depend on this number. The ability to predict intelligence reached statistical significance at 16 time frames for the highest maxima and at 24 time frames for the lowest minima (Fig. 4e,f). This result shows that individual intelligence scores can be predicted from as few as 16 time frames, equal to < 1.51% of a 10 min resting-state fMRI session (TR = 645 ms, 860 TFs), as long as temporal independence of the selected time frames is ensured. These are more time frames than necessary to reconstruct the static functional connectome (e.g., MxCo contained less than 11 frames), indicating that more independent data points are required to predict a complex human trait like general intelligence.

### 3.5. Randomly selected time frames predict intelligence as good as time frames of maximal cofluctuation

As states of highest maxima (Mx) are not significantly superior to states of lowest minima (Mn) in predicting intelligence (Fig. 4b,c), we next asked whether comparable prediction performances could be obtained when using an equal number of time frames randomly sampled from a uniform distribution over all time frames. Similar to the original HiCo and LoCo selections, also highest maxima (Mx) significantly outperformed the randomly selected set of time frames in reconstructing the static connectivity matrix, whereas the lowest minima (Mn) performed significantly worse (orange and green line in comparison to the gray area in Fig. 4d). In contrast, when predicting intelligence, the random sets of time frames performed just as good as sets with an equal number of highest maxima and lowest minima (Fig. 4e,f) but significantly worse than an equal number of HiCo and LoCo frames (Fig. S5). This suggests that while reconstruction similarity to the static FC primarily depends on the strength of cofluctuation, the prediction of intelligence is mainly determined by the number independent time frames.

### 3.6. Intelligence prediction involves multiple functional brain networks

The analyses reported so far used information from all possible functional brain connections for predicting intelligence. However, the brain can be decomposed into multiple non-overlapping networks (or modules) associated with different cognitive functions (e.g., Fox et al., 2005). To assess their relative contributions to the prediction of intelligence, we repeated our analyses for a) static connectivity (all time frames) and b) the 43 highest maxima, considering seven established functional brain networks (Yeo et al., 2011). Specifically, we analyzed, in a first step, the prediction performance of only the connections within a network and of those between a specific pair of networks. As illustrated in Fig. 5a, static connections within the visual network significantly predicted intelligence (*p* < .05, Bonferroni corrected for 28 comparisons, i.e., 21 between-network analyses and 7 within-network analyses). When using the set of 43 highest maxima, only connections between the default-mode network and the limbic system (Fig. 5b) could significantly predict intelligence (*p* < .05, Bonferroni corrected for 28 comparisons). Secondly, we further explored the relevance of the seven functional networks for prediction of intelligence by analyzing the change in overall prediction performance when removing all connections, a specific network was involved in. No significant changes in prediction performances were observed when removing one network from static connectivity, while removing the fronto-parietal network for cognitive control from the selection of 43 highest maxima reduced the prediction performance significantly, however, only when not correcting for multiple comparisons (Fig. 5d; *p* < .05; both analyses Bonferroni corrected for seven repeated analyses, i.e., one per network). In sum, these results suggest that multiple functional brain systems contribute to the prediction of intelligence.

**Fig. 5.**
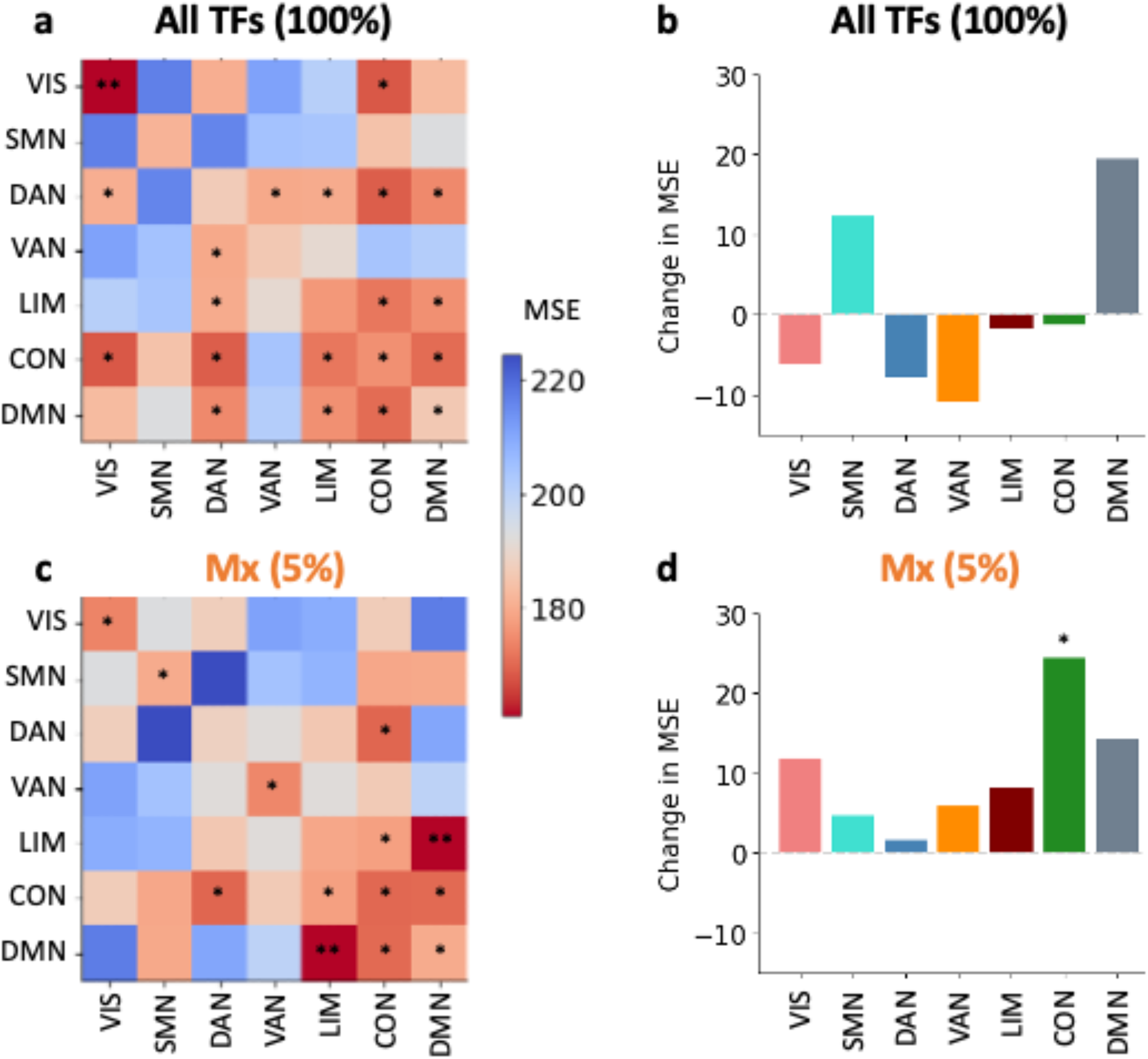
Multiple functional brain networks contribute to the prediction of intelligence. Intelligence (FSIQ; WASI, Wechsler, 1999) was predicted with CMEP from (**a, b**) static functional connectivity (all time frames; TFs) and (**c, d**) from the 43 highest maxima of the global cofluctuation time series (Fig. 1e). In (**a, c**) prediction performance (mean squared error; MSE) of connectivity within or between seven functional brain networks (Yeo et al., 2011) was analyzed by selecting only the specific within or between network connections, while (**b, d**) illustrates the change in predictive performance (MSE) after removing all connections a respective network was involved in. Significance was determined by a non-parametric permutation test with 1,000 iterations. * if *p* < .05 uncorrected for multiple comparisons and ** if *p* < .05 Bonferroni corrected for multiple comparisons (28 comparisons, *p* < .0018 in **a** and **c** and seven comparisons, *p* < .007 in **b** and **d**). VIS, visual network; SMN, somatomotor network; DAN, dorsal attention network, VAN, ventral attention network; LIM, limbic network; CON, control network; DMN, default mode network.

### 3.6. Robustness control analyses

Multiple control analyses were performed to further evaluate the robustness of our findings. As the choice of the cross-validation strategy can potentially influence the results of predictive modelling approaches (Varoquaux, 2018), we first repeated our analyses for static functional connectivity and the 43 highest maxima using stratified 10-fold cross validation instead of LOO cross validation and obtained highly similar results (Tab. S1). Given that both the cognitive ability measure used in this study (FSIQ) and functional brain connectivity may vary with age (e.g., Deary et al., 2009; Geerligs et al., 2015) we secondly repeated our analyses using age-adjusted intelligence scores operationalized as residuals resulting from a linear regression predicting the mean FD adjusted FSIQ score (ўa) from age (*x*). As illustrated in Tab. S1, these analyses resulted in overall comparable findings for static connectivity and 43 highest maxima. Third, as functional connectivity estimates can be critically influenced by motion (e.g., Ciric et al., 2017; Power et al., 2012), we investigated whether there exists a temporal correspondence between the 43 highest maxima and high-motion time frames. This was not the case (Fig. S6a). Fourth, we tested whether the temporal distribution of the 43 highest maxima itself might be associated with variations in intelligence and observed that this also was not the case (*r* = .02, *p* = .78; Fig. S6b). In sum, the control analyses support the robustness of our findings.

### 3.7. External replication

Even though all of the results outlined above were thoroughly cross validated and tested for different confounding effects, potential remaining influences of sample-specific characteristics can only be ruled out by external replication (Cwiek et al., 2021). We therefore repeated our analyses using the four resting-state fMRI scans from the Human Connectome Project (HCP; *N* = 831, age range: 22 – 36 years; 100 nodes, Schaefer et al., 2018; see Methods for details) each cropped to the same length as the time series of the primary sample (860 TFs). Given that the HCP data do not provide a full-scale IQ measure, general cognitive ability was operationalized as latent *g*-factor (Spearman, 1904) computed from 12 cognitive performance scores (following Dubois et al., 2018; Fig. S1). This second dataset corroborated our prediction results (Tab. S2; Fig. S7): Static functional connectivity derived from all time frames could significantly predict intelligence (averaged across the four scans: *r* = .23, *p* < .001), while highest and lowest cofluctuation states could not (HiCo: *r* = .08, *p* > .05; LoCo: *r* = .02, *p* > .05). As in the primary dataset, restricting the selection to the maxima/minima during these highest/lowest cofluctuation states (< 17 TFs) resulted in only a slight decrease in reconstruction similarity (Tab. S2 in comparison to Tab. 1) and prediction performance (MxCo: *r* = .07, *p* > .05; MnCo: *r* = -.03, *p* > .05). Finally, also in the replication sample, the 43 highest maxima and 43 lowest minima allowed for significant prediction of intelligence (Mx: *r* = .23, *p* < .001; Mn: *r* = .18, *p* < .05). Note that the reconstruction similarity to static connectivity was again highest in the latter cases (Mx: *r* = .97, Mn: *r* = .74). Finally, comparisons with randomly selected time frames and across increasing numbers of time frames resulted in similar findings as reported above for the original analyses (Fig. S7c,d).

Network-specific analyses in the replication sample yielded qualitatively similar findings, i.e., significant predictive power of connections linking the default mode, the fronto-parietal, and the attention networks (Fig. S8a,b). Also, in no case prediction performance (MSE) decreased significantly when removing one network (all *p* > .05; Fig. S8c,d). While lacking the specificity observed in the primary sample, this supports the conclusion that multiple networks contribute to the prediction of intelligence. Lastly, an additional control analysis applying the 114-nodes Yeo-atlas (Yeo et al., 2011) to the replication sample yielded similar results and, thus, demonstrated the robustness of findings also against variations in different node parcellation schemes (Tab. S3). Overall, the results of the external replication support the generalizability of our findings to different samples, different scanning and preprocessing parameters, different age cohorts, and to different measures of general cognitive ability.

## 4. Discussion

Recent work has shown that brief states of particularly high functional connectivity reflect fundamental properties of individuals’ functional connectomes measured across several minutes of resting-state fMRI (Betzel et al., 2022a; Esfahlani et al., 2020). Here, we explored whether these network-defining and person-specific states of high brain-wide cofluctuation – as well as other network states – carry information about individual differences in general cognitive ability (intelligence). We first replicated the basic phenomenon that the fundamental structure of static functional connectivity is driven by a small number of such high cofluctuation states (Esfahlani et al., 2020). Further a machine learning-based prediction framework (CMEP) was developed to predict intelligence from functional connectivity created from only such states. However, neither the high cofluctuation states nor the states of particularly low cofluctuation were predictive for intelligence. This low prediction performance of high cofluctuation states as defined by Esfahlani et al. (2020) and Betzel et al., (2022a) potentially results from the high number of temporally adjacent and thus correlated frames that carry little independent information. Secondly, we reveal that intelligence can be predicted from equally small selections of fMRI time frames – when such time frames are temporally independent. This holds true for selections of maxima and minima within the cofluctuation time series, as well as for random selections of time frames. Lastly, we show that intelligence can be predicted from selections of as few as 16 time frames, and that intelligence prediction relies on multiple functional brain networks, including the visual, the attention, the limbic, the fronto-parietal control, and the default mode systems. The replication of all results in an independent sample suggests generalizability of our findings to different populations, processing pipelines, and to various measures of cognitive ability.

### 4.1. How much brain data is required to predict human cognition?

A large number of recent studies demonstrated that cognitive abilities can be predicted from functional brain connectivity measured with fMRI (e.g., Dhamala et al., 2021; Finn et al., 2015; Thiele et al., 2022). The recent advent of time-resolved brain connectivity analyses (Airan et al., 2016; Esfahlani et al., 2020) and of time-varying connectivity approaches in general (for review see Lurie et al., 2020) has made it possible to investigate the relationship between specific states of functional connectivity and cognition in more detail. Previous work has strongly focused on time frames of particularly high cofluctuation and demonstrated their ability to capture idiosyncratic information (Betzel et al., 2022a; Cutts et al., 2023; Sporns et al., 2021). In fact, our results suggest that the ability to predict intelligence is rather independent from the strength of cofluctuation but critically depends on the availability of sufficient independent data.

Further, our finding that prediction performance of intelligence increases with the number of time frames and that sufficient temporal independence is required, contradicts earlier studies proposing that scans as short as three to four minutes are sufficient to characterize individual subjects comprehensively (Airan et al., 2016; Byrge & Kennedy, 2019) and that even less than two minutes of resting-state fMRI can be used to build robust individual connectotypes (i.e., idiosyncratic connectivity properties; Miranda-Dominguez et al., 2014). In such short scanning durations, it might not be possible to detect a sufficient number of frames that contain enough independent trait-relevant information. On the one hand, our findings support the benefit of analyzing human brain connectivity with temporally fine-grained methods that prevent the temporal averaging step (see also O’Connor et al., 2022). On the other hand, we argue towards the relevance of longer scanning durations for neuroimaging research on individual differences in complex human traits to ensure capturing sufficient trait-relevant connectivity states.

### 4.2. Which brain states reflect individual differences in intelligence?

The selection of only few time frames provides a method to clarify whether or not individual differences in intelligence depend upon specific brain states - as recently proposed by the Network Neuroscience Theory of Intelligence (NNT; Barbey, 2018). We therefore defined a brain connectivity state as a small selection of time frames from the fMRI connectivity time series that are all characterized by specific connectivity properties, and we probed such selections for their ability to predict individual intelligence scores.

The successful recovery of individual-specific connectomes from a limited number of highest connectivity brain states (Esfahlani et al., 2020) made these states a plausible first target. Also, Ladwig et al. (2022) suggested that focusing on time frames of higher cofluctuation increases the signal to noise ratio and thus increases the probability of detecting trait-relevant information. In contrast, in our study time frame selections with higher cofluctuation did not differ significantly from selections with lower cofluctuation regarding their predictive ability. This result is supported by Sasse et al. (2022), who find that lower and intermediate cofluctuation time frames provide higher subject specificity as well as highest phenotype predictions. Ladwig et al. (2022) further demonstrated that high cofluctuation states are not necessarily special: removing these states from the connectivity time series did not impair the ability to derive individual connectomes, and random selections were equally capable of reconstructing static connectivity matrices. Relatedly, our result that even randomly selected time frames can predict intelligence as good as highest cofluctuation maxima suggests that intelligence-relevant information is not reflected in a specific brain-network organizational state (such as the high cofluctuation states) but is rather present in many fMRI time frames distributed across the entire scan. This assumption is further supported by the observation that the selection of time frames that allowed for highest predictive performance (i.e., Mx, Mn) includes a higher number of distinct spatial coactivation patterns and may thus depict more independent information. Our study adds to the large literature on the occurrence of individual brain states (Allen et al., 2014; Liu et al., 2018; Medaglia et al., 2018) but additionally analyzes such states on a frame-by-frame level as well as regarding their relevance for cognitive abilities. To summarize, intelligence does not seem to be characterized by a specific brain state, but rather by the ability of the human brain to explore different brain states over time.

### 4.3. Implications for understanding the brain bases of intelligence

The involvement of multiple functional brain networks in the prediction of intelligence from functional connectivity, observed here, is well in line with established neurocognitive theories of human intelligence (Parieto-Frontal Integration Theory, P-FIT, Jung & Haier, 2007; Multiple Demand System, MD, Duncan, 2010) and more recent meta-analytic findings (Basten et al., 2015) suggesting that a widely distributed system of brain regions is implicated in individual differences in intelligence. Further support for the relevance of whole-brain analyses comes from a more theoretical perspective claiming that cognitively highly demanding activities – like complex problem solving – require the integration of information that is distributed widely across the brain (Duncan et al., 2020). We observed consistent relevance of the default-mode and the fronto-parietal networks in the prediction of intelligence from the full time series as well as the selection of 43 highest maxima, corroborating previous results that these networks are especially relevant for individual differences in intelligence (e.g., Dubois et al., 2018; Finn et al., 2015; Thiele et al., 2022; for review see Hilger & Sporns, 2021). The reduction in predictive power of connections within the dorsal attention and visual network when comparing predictions based on all time frames to those based on the highest maxima may potentially result from the underrepresentation of the inter- and intra-connectivity of smaller networks (and thus the overrepresentation of larger networks) when computing the whole-brain cofluctuation (Betzel, et al., 2022b). However, best predictions were achieved with whole-brain connectivity, highlighting the positive effect of information from all networks on predicting intelligence.

### 4.4. Introducing CMEP – a data-driven prediction framework for neuroscience

Finally, the here-introduced prediction framework - covariance maximizing eigenvector-based predictive modelling (CMEP) - has some notable advantages in comparison to previously used prediction methods for functional neuroimaging data. First, we demonstrated in multiple control analyses superior robustness of prediction results derived from CMEP in comparison to those derived from the to date most frequently used approach in network neuroscience (CPM; Finn et al., 2015; Shen et al., 2017). Second, CMEP is a completely data-driven framework, thereby making it unnecessary to (arbitrarily) select a threshold parameter (e.g., to identify most relevant brain connections). This reduces the researchers’ degrees of freedom and thereby facilitates the reproducibility of results (Gilmore et al., 2017; Poldrack, 2019). We therefore propose that CMEP is better suited to extract trait-like connectivity characteristics. However, compared to CPM, CMEP comes with slightly higher computational costs (1.14 x longer training time in comparison to CPM) and although the generated eigenvectors allow for more detailed functional interpretations (Fig. 2c), they produce more complex features than the two network averages used in CPM (i.e., a positive network and a negative network; Finn et al., 2015; Shen et al., 2017). Note however, that all tested models for intelligence prediction seem to be not transferrable to a completely new data set - an observation which is most likely caused by differences in sample characteristics, data acquisition parameters, preprocessing strategy, and in the predicted cognitive measure. Despite these limitations, our results suggest CMEP as a promising candidate for broad applications in neuroscientific studies testing for brain-behavior relationships.

## 5. Limitations and Future Directions

Our results are in line with the recent proposal in Barbey (2018) that different brain states underly human cognition, however, we did not probe for the dynamic reconfiguration between such states. We have, however, recently demonstrated that persons with higher IQ scores show higher stability of network modularity over time – an effect that was in part driven by the same functional systems as identified here (Hilger, et al., 2020a). These former results together with our current observation that even random selections of time frames allow for significant prediction of intelligence contradicts the NNT’s proposal that the brain’s ability to flexibly access a specific set of network states is essential for higher levels of intelligence (Barbey, 2018). Whether higher network stability, higher network flexibility or neither of the two facilitates human cognition requires clarification in future work. However, our former study (Hilger, et al., 2020a) and the NNT (Barbey, 2018) imply that individual differences in intelligence relate to the dynamic reconfigurations of functional connectivity over time, while our current study adds that the most predictive brain connectivity states include more heterogeneous coactivation patterns that can only be captured by ensuring sufficient temporally independent fMRI data frames. However, as this requires sufficiently long scan durations, such types of analyses cannot directly be transferred to all standard fMRI settings. In sum, this research highlights the need for time-varying connectivity analyses and sufficient long scan durations.

Our study focuses on the analysis of distributed states of functional MRI during rest. Recent literature has also highlighted the potential benefits of including anatomic MRI as well as diffusion tensor imaging (DTI) into prediction models (e.g., Dhamala et al., 2021; Litwińczuk et al., 2022; Ooi et al., 2022; Popp et al., 2023; Sarwar et al., 2021). Future work should extend this research and simultaneously include a broad range of structural parameters such as anatomical connectivity combined with cortical thickness, cortical surface area, or gyrification. Additionally, task-induced functional connectivity has been related to individual differences in cognitive ability (e.g., Greene et al., 2018; Fong et al., 2019; Jiang et al., 2020) and represents together with dynamic fMRI analyzes another promising future perspective (e.g., Chen et al., 2022).

The prediction performances in our study were in a comparable range to previous large-scale studies predicting intelligence or other complex human traits from static brain connectivity including all time frames (He et al., 2020; for review see Dizaji et al., 2021) or even from brain structure (Hilger, et al., 2020b; mean absolute error ∼10 IQ points). However, it has recently been proposed that the investigation of brain-behavior relations requires large sample sizes (DeYoung et al., 2022; Marek et al., 2022; Rosenberg & Finn, 2022) and that replication studies observe small effect sizes (Marek and colleagues (2022) found the largest 1% of replicable univariate effects to be between |*r*| = .06 and .16). Despite limited sample sizes in the range of hundreds our study demonstrates (via external validation) that the combination of cross-validation and sophisticated analyses approaches like CMEP allows to reliably identify brain-behavior associations - a promising direction for future research.

## 6. Conclusion

We introduced an eigenvector-based prediction framework to show that functional connectivity estimated from as few as 16 temporally separated time frames allow to significantly predict individual differences in general cognitive ability (*N* = 263). In contrast and against previous expectations, we revealed that network-defining time frames of particularly high or low cofluctuation are not predictive of intelligence, such that much more frames would be required to achieve a predictive performance comparable to the complete time series. Further we showed that multiple functional brain networks contribute to the prediction of cognitive ability from small selections of the fMRI time series, and all results replicate in an independent sample (*N* = 831). Overall, our results reveal that whereas relatively few time frames of brain connectivity are sufficient to derive fundamentals of person-specific functional connectomes, temporally distributed information and hence ultimately more data is necessary to extract information about cognitive abilities from functional connectivity time series. Importantly and in contrast to recent proposals, intelligence-predictive information is not restricted to specific brain connectivity states, but rather reflected within many different brain states that are temporally widely distributed over the whole brain connectivity time series. To detect such states requires sufficient amounts of neuroimaging data, making longer scanning durations essential for reliable prediction of complex human traits.

## Declaration of Competing Interest

Declaration of interest: none.

## CRediT authorship contribution statement

**Maren Wehrheim:** Conceptualization, Methodology, Formal analysis, Writing – Original Draft, Visualization. **Joshua Faskowitz:** Methodology, Data curation; Writing – Review & Editing. **Olaf Sporns:** Methodology, Writing – Review & Editing. **Christian Fiebach:** Conceptualization, Resources, Writing – Review & Editing; Funding acquisition. **Matthias Kaschube**: Conceptualization, Methodology, Resources, Writing – Review & Editing, Supervision; Funding acquisition. **Kirsten Hilger:** Conceptualization, Methodology, Resources, Writing – Original Draft, Supervision; Funding acquisition.

## Acknowledgements

The authors thank the Nathan S. Kline Institute for Psychiatric Research (NKI; Nooner et al., 2012), founded and operated by the New York State office of mental health, for providing the primary data set for the current study, and the Human Connectome Project (Van Essen et al., 2013) for providing data of the replication sample. Finally, the authors thank Makoto Fukushima for providing the preprocessed NKI data.

## Funding

The research leading to these results has received funding from the German Research Foundation (DFG Grant FI 848/6-1), the European Community’s Seventh Framework Programme (FP7/2013) under Grant agreement n° 617891, and the Alfons and Gertrud Kassel-Stiftung (MW, CF, MK). In part, the project was also supported by Lilly Endowment, Inc., through its support for the Indiana University Pervasive Technology Institute. KH received funding from the German Research Foundation (DFG Grant HI 2185/1-1).

## Supplementary Material

Supplementary Material associated with this article can be found in a separate document.

## Notes

https://github.com/kaschube-lab/CMEP

https://fcon_1000.projects.nitrc.org/indi/enhanced/

https://www.humanconnectome.org/study/hcp-young-adult

https://github.com/faskowit/app-fmri-2-mat

https://fmriprep.readthedocs.io/en/20.2.5/workflows.html

### Competing Interest Statement

The authors have declared no competing interest.

### Summary of Updates

Figure 1 and 4 revised; Section on HCP data clearified; Additional section in discussion

