## Supplementary material for "Few temporally distributed brain connectivity states predict human cognitive abilities": SI

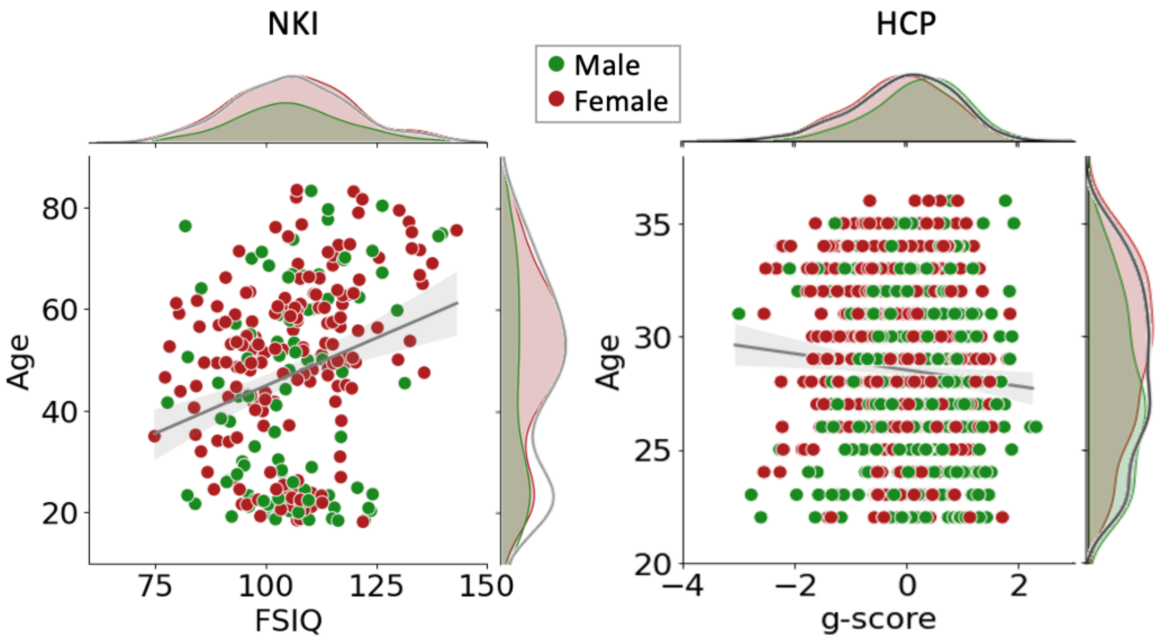


**Supplementary Fig. S1.** Distribution of intelligence test scores (FSIQ in the main sample, NKI, Nooner et al., 2012, and latent *g*-factor in the replication sample, HCP, Van Essen et al., 2013) and age in both data sets. In the main sample (NKI), intelligence test scores were weakly correlated with age ($r = .27$; $p < .001$) with a mean (standard deviation) of 101.78 (13.14) and 47.14 (18.25) for intelligence and age, respectively. Within the male and female subgroup, the test scores and age were correlated with $r = .18$ ($p = .09$) and $r = .27$ ($p < .001$), respectively. Intelligence tests scores had a mean of 101.59 (101.87) and standard deviation of 12.56 (13.44) for the male (female) subgroup. Age was less similarly distributed between the gender subgroups with a mean of 42.93 (49.37) and a standard deviation of 20.07 (16.78) for the male (female) subgroup. Right: Age and *g*-score were weakly negatively correlated in the replication sample (HCP: $r = -.09$; $p = .014$). For the *g*-scores and age we observed a mean (standard deviation) of 0 (.89) and 28.55 (3.72), respectively. Within the male and female subgroup, intelligence and age were correlated with $r = .01$ ($p = .78$) and $r = -.09$ ($p = .06$), respectively. The *g*-score showed a mean of .19 (-.17) and standard deviation of .86 (.87) for the male (female) subgroup. For age we observed a mean of 27.63 (29.36) and standard deviation of 3.63 (3.61) for the male (female) subgroup.

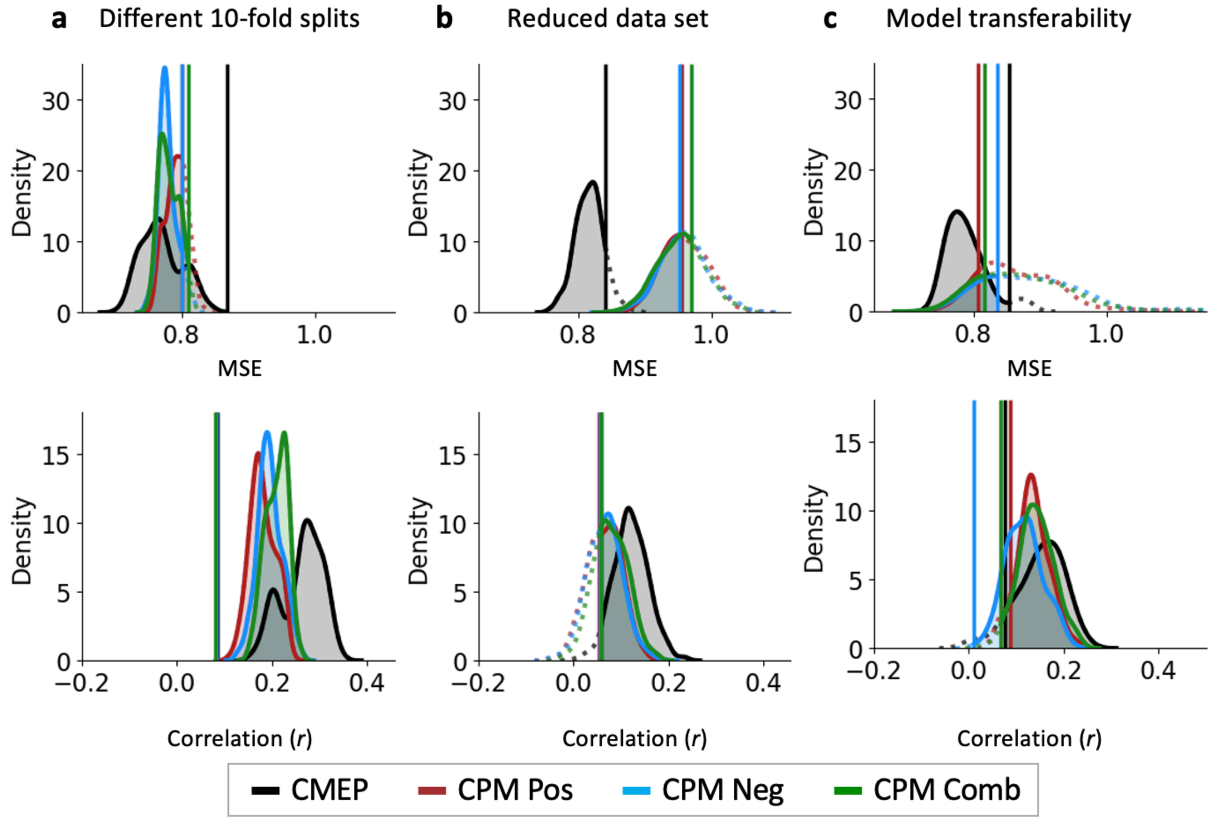


**Supplementary Fig. S2.** Superior prediction robustness of covariance maximizing eigenvector-based predictive modelling (CMEP) relative to connectome-based predictive modelling (CPM) in the replication sample. Prediction performances (mean squared error, MSE, and Pearson correlation between observed and predicted intelligence scores, between observed and predicted intelligence scores) from static (time-averaged) connectivity were compared via three validity analyses between CMEP and CPM (Finn et al., 2015; Shen et al., 2017). All analyses were conducted for CMEP (black, all brain connections), and three CPM prediction pipelines based on positive connections (red), negative connections (blue), and a combination of both (green, all connections). (a) Robustness across different data set splits. Data were randomly (100 times) split into 10 folds for cross validation. (b) Robustness across different sample sizes. Within stratified 10-fold cross-validation, the training sample was randomly (100 times) reduced to 10% of the original test-sample size. (c) Transferability of the models to a new data set. Models were trained on the replication sample (HCP) and tested on the primary sample (NKI). Both samples were parcellated into the 200 nodes schemata (Schaefer et al., 2018) and all intelligence scores were first standardized and then after prediction mapped back to the original scale for better comparability. The training data were randomly bootstrapped (100 times) to account for different compositions of the training data set. The vertical solid lines indicate the significance threshold ($p$ < .05) for each model. Models that were found to be significant are indicated by a shaded area and solid line, insignificant models are depicted with dotted lines.

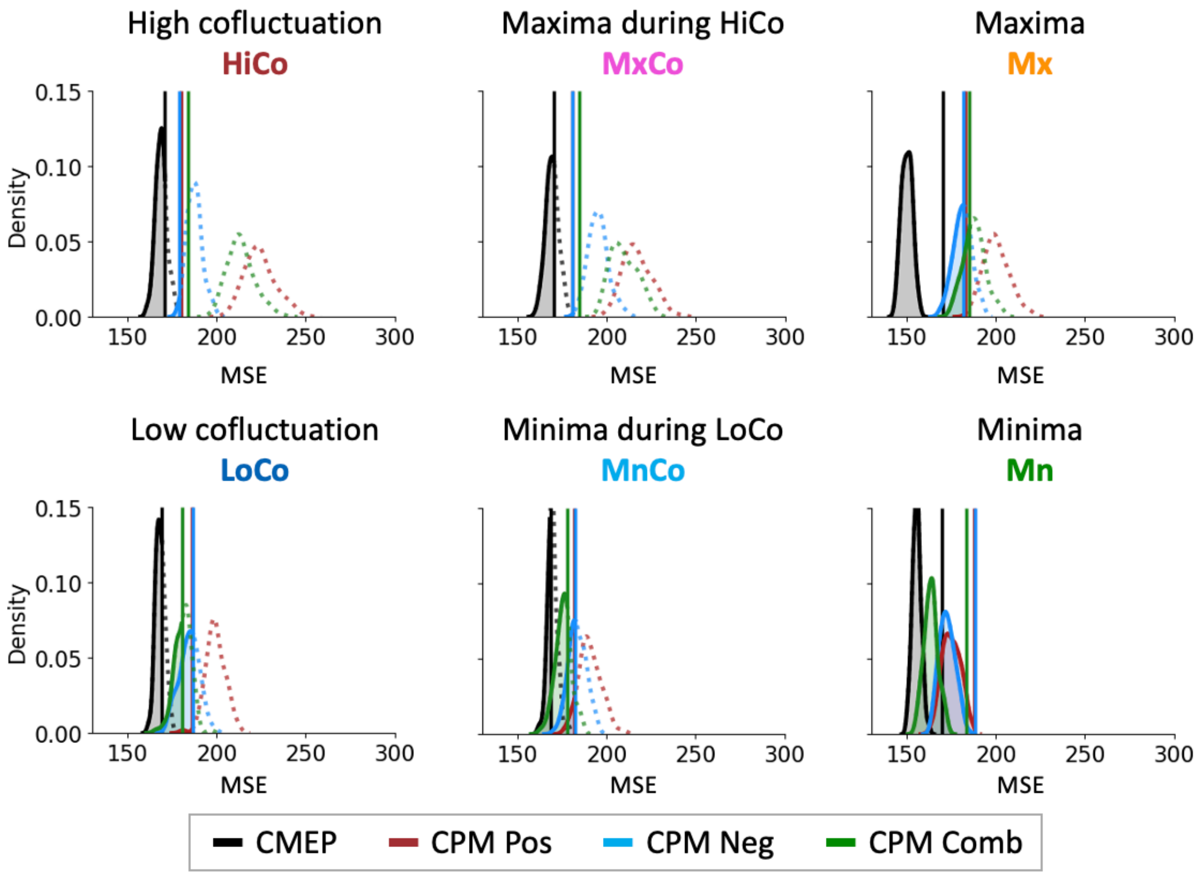


**Supplementary Fig. S3.** Prediction robustness of Covariance Maximizing Eigenvector-Based Predictive Modelling (CMEP) in contrast to connectome-based predictive modelling (CPM) for all six different connectivity states. Prediction results are evaluated with the mean squared error (MSE) between predicted and observed intelligence scores (FSIQ; WASI, Wechsler, 1999). Prediction features were derived from one of six different connectivity states (see Fig. 1). Robustness is operationalized as the empirical distribution of prediction performances resulting from 100 different cross-validations splits (10-fold). CMEP (black) is implemented as described in the Methods section and illustrated in Fig. 2. In CPM (Finn et al., 2015; Shen et al., 2017) positive and negative connectivity strengths are calculated as the sum over all functional connections that are significantly positively (red) or negatively (blue) correlated with intelligence above a given threshold (here: $p < .001$). These positive and negative connectivity strengths serve separately as features to fit a linear regression model to predict intelligence. Note that as CMEP does not differentiate between positive and negative functional brain connectivity, we additionally fitted CPM with positive *and* negative connectivity strengths (CPM Comb, green). The vertical solid lines indicate the significance threshold ($p < .05$) for each model. Models that were found to be significant are indicated by a shaded area and solid line, insignificant models are depicted with dotted lines.

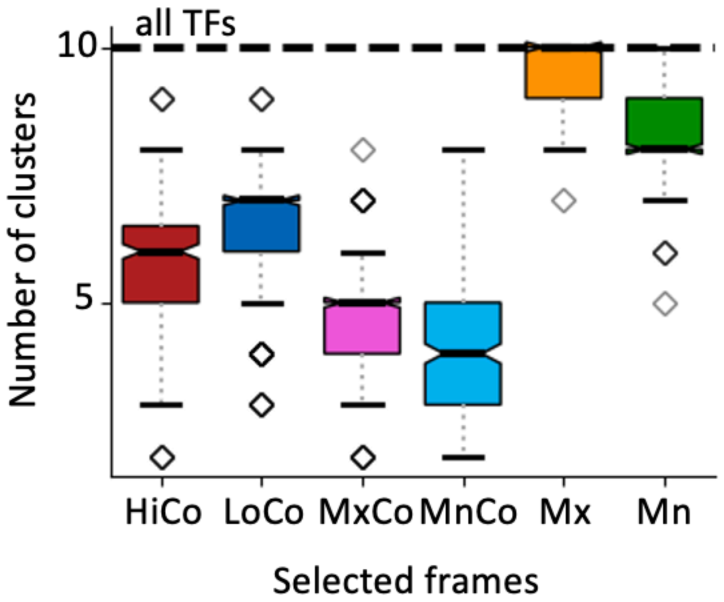


**Supplementary Fig. S4.** Temporally distributed time frames as depicted in cofluctuation maxima and minima (Mx, Mn) engage more spatially separable coactivation patterns than temporally adjacent states of highest/lowest cofluctuation (HiCo, LoCo). Following the literature on coactivation patterns (CAPs, Liu et al., 2018) the fMRI activation time series was divided into ten different clusters using the k-means clustering algorithm. We report the number of clusters (coactivation patterns) that were engaged in each of the six different time frame selections (see Fig. 1a). Note that all results are highly similar also when using k = 5,…, 20 clusters.

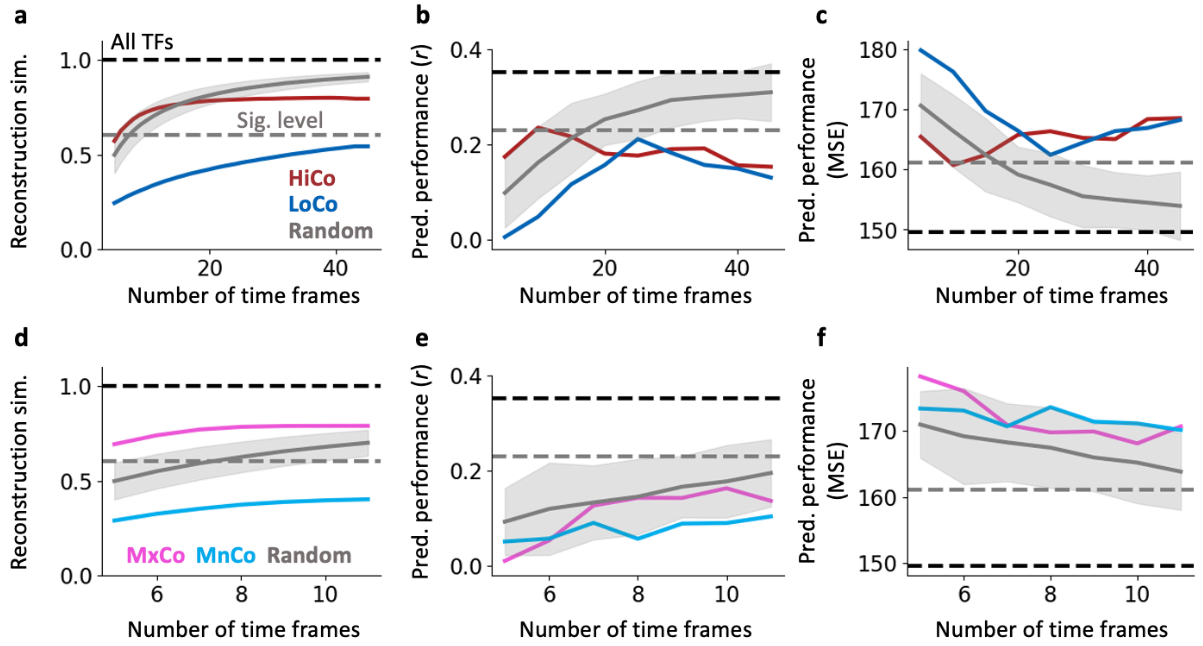


**Supplementary Fig. S5.** Reconstruction similarity and prediction performance for different numbers of time frames. Reconstruction similarity (left; pearson correlation *r* between reconstructed connectivity and static functional connectivity; dashed lines) and prediction performance (center: correlation between predicted and observed scores, *r*; right: mean squared error, MSE) across 100 different 10-fold cross validation splits. (**a-c**): Functional connectivity was reconstructed from time frames with decreasing strength of cofluctuations (as indicated by RSS values) starting with the five time frames of highest cofluctuation (red) and from time frames with increasing strength of cofluctuations starting with the five time frames of lowest cofluctuation (blue). (**d-f**): Functional connectivity was reconstructed from only the maxima/minima within the highest/lowest cofluctuation time series (pink/light blue). For further comparability a null model (gray) was generated from 100 random time frames that were uniformly selected. The translucent band around the mean prediction performance of the random selections indicates the standard deviation of prediction results.

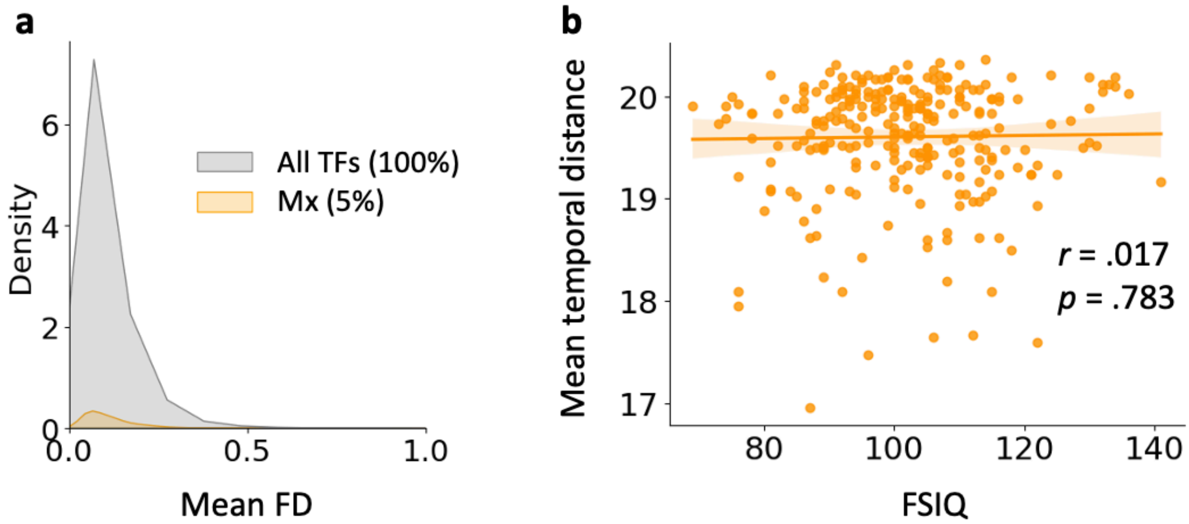


**Supplementary Fig. S6.** Control analyses results. (**a**) Relative independence of the 43 highest maxima connectivity states from time frames of high head motion. Empirical distribution of in-scanner head motion (operationalized as mean framewise displacement) for the complete time series (gray) and the 43 highest maxima connectivity states only (orange). (**b**) Associations between intelligence (FSIQ; WASI, Wechsler, 1999) and the mean temporal distribution of 43 highest maxima connectivity states. Each dot represents one subject, and the best-fit regression line is highlighted with a translucent band corresponding to the 95%-confidence interval. *r*, Pearson correlation coefficient, *p*, 2-tailed *p*-value.

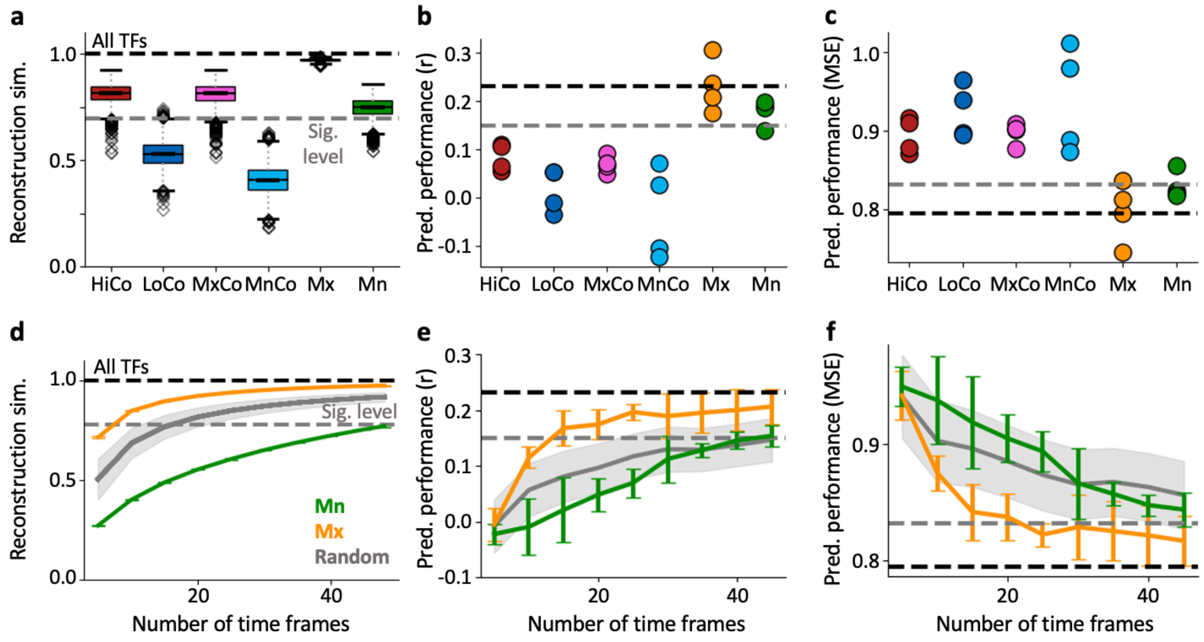


**Supplementary Fig. S7.** The performance to predict intelligence depends on the number of temporally independent time frames rather than on reconstruction similarity also in the replication sample (HCP). (**a**) Reconstruction similarity of six different connectivity states operationalized as Pearson correlation between static functional connectivity (constructed from all time frames; TFs) and connectivity matrices reconstructed from six different selections of TFs. Boxplots depict the mean and quartiles of the subject-specific reconstruction similarity for all different connectivity states and across all four scans. The whiskers show the 1.5 x interquartile ranges. Outliers are represented by diamonds. Performance to predict intelligence (*g*-score) for the six different connectivity types from using the CMEP prediction framework (see Fig. 2, (**b**) correlation between predicted and observed scores, *r*; (**c**) mean squared error, MSE). Each dot represents one scan session. (**d**) Reconstruction similarity and performance (correlation, **e**; MSE, **f**) to predict intelligence as a function of the number of (randomly selected) time frames comprising cofluctuation maxima or cofluctuation minima (orange or green dots in Fig. 1e). Gray lines represent reconstruction similarity (**d**) and predictive performance (**e**) and (**f**) from randomly selected time frames (see Methods for further details about the null model). Orange and green lines represent results from highest maxima and lowest minima connectivity states averaged across scans. The whiskers represent the standard deviation across scans. Note that for prediction performances only the two cases (highest maxima and lowest minima) are illustrated that allow for significant prediction of intelligence, i.e., 43 highest maxima, Mx; 43 lowest minima, Mn. The upper bounds (black dashed lines) represent reconstruction similarity (**a, d**) or prediction performance (**b, c, e, f**) using all TFs. The lower grey dashed line reflects the approximate 5% significance level (determined as average over all seven model's significance levels) of the within-subject similarity of static functional connectivity (**a, d**) or intelligence prediction performance (**b, c, e, f**, see Methods). HiCo, high cofluctuations; LoCo, low cofluctuations; MxCo, maxima during HiCo; MnCo, minima during LoCo; Mx, Maxima; Mn, Minima (see also Fig. 1e).

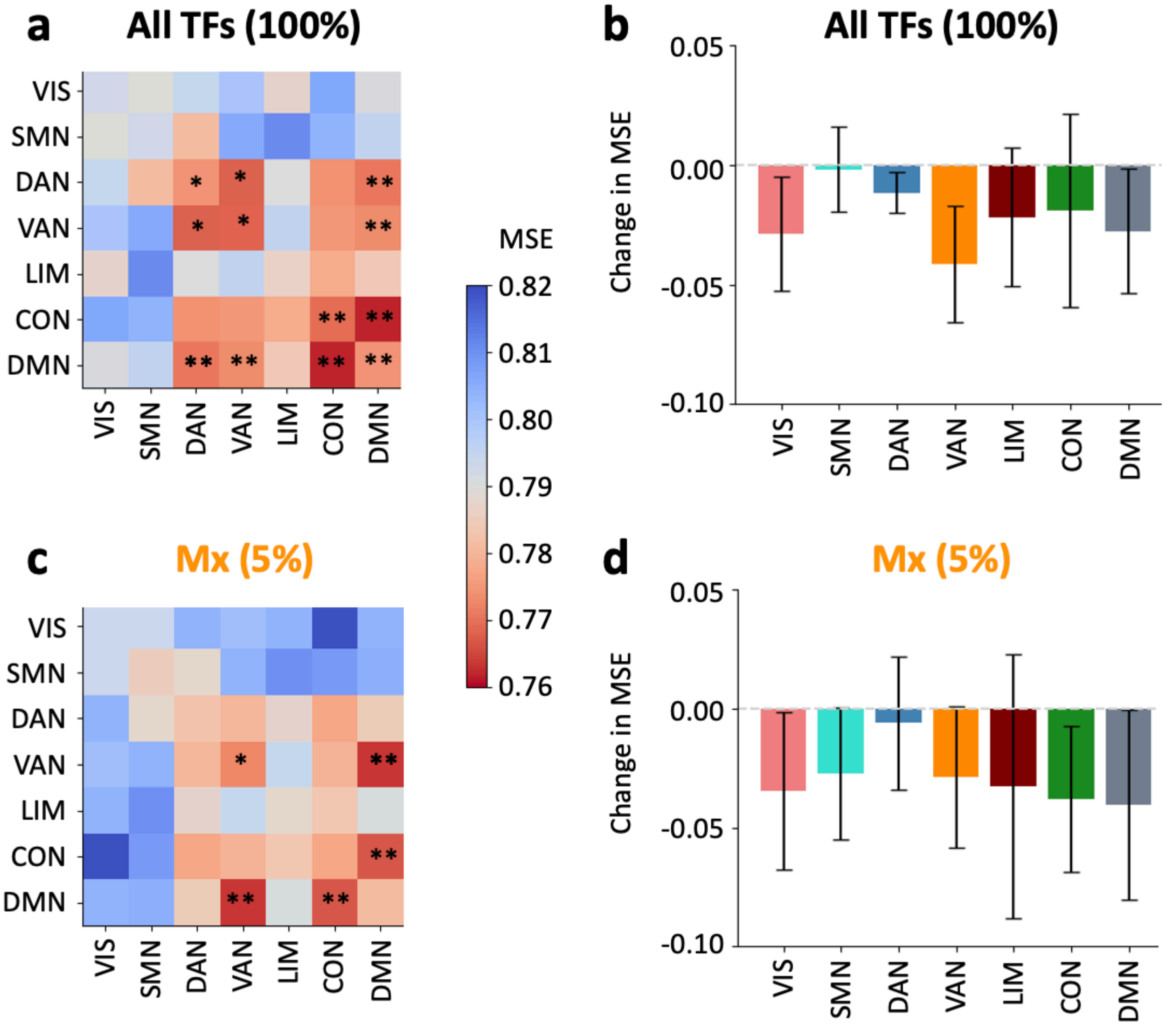


**Supplementary Fig. S8.** Multiple functional brain systems contribute to the prediction of intelligence also in the replication sample (HCP). Intelligence (*g*-score) was predicted with CMEP from (**a, b**) static functional connectivity (all time frames; TFs) and (**c, d**) from the 43 highest maxima of the global cofluctuation (Fig. 1e). In (**a, c**) prediction performance (mean squared error; MSE) of connectivity within or between seven functional brain networks (Yeo et al., 2011) was analyzed by selecting only the specific within or between network connections, while (**b, d**) illustrates the change in prediction performance (MSE) after removing all connections a respective network was involved in. All results are depicted as the mean across the four scan sessions and whiskers indicate the standard deviation across the sessions. Significance was determined by a non-parametric permutation test with 1,000 iterations. * if $p < .05$ uncorrected for multiple comparisons and ** if $p < .05$ Bonferroni corrected for multiple comparisons (28 comparisons, $p < .0018$ in **a** and **c** and seven comparisons, $p < .007$ in **b** and **d**). VIS, visual network; SMN, somatomotor network; DAN, dorsal attention network, VAN, ventral attention network; LIM, limbic network; CON, control network; DMN, default mode network.

**Supplementary Tab. S1**

Prediction results for 10-fold cross validation instead of leave-one-out (LOO) and when controlling intelligence scores for potential age effects

| | **10-fold cross validation** | | **Age-adjusted intelligence scores** | |
|---|---|---|---|---|
| | Static functional connectivity (All TFs) | Maxima (Mx) | Static functional connectivity (All TFs) | Maxima (Mx) |
| *r* | .35** | .37** | .32** | .36** |
| MSE | 149.59** | 147.74** | 142.05** | 138.11** |
| RMSE | 12.23** | 12.16** | 11.92** | 11.75** |
| MAE | 9.59** | 9.62** | 9.48* | 9.37** |

*Note*: Results are listed for static functional connectivity (All TFs) and general maxima connectivity states (Mx). Model performance metrics reflecting the fit (*r*) or error (MSE, RMSE, MAE) between predicted and observed intelligence scores: Pearson correlation coefficient (*r*), mean squared error (MSE), root mean squared error (RMSE), and mean absolute error (MAE). Significance is determined by a non-parametric permutation test with 1,000 iterations and indicated as ** if $p < .001$, * if $p < .05$.

**Supplementary Tab. S2**

Prediction of intelligence from functional brain connectivity for the replication sample using the Schaefer 100 nodes partition (Schaefer et al., 2018)

| | TFs | Reconstruction similarity | *r* | MSE | RMSE | MAE |
|---|---|---|---|---|---|---|
| Static functional connectivity | 860 | n/a | .23** | 0.79** | 0.89** | 0.71** |
| Highest cofluctuations (HiCo) | 43 | .81 | .08 | 0.89 | 0.95 | 0.75 |
| Lowest cofluctuations (LoCo) | 43 | .53 | .02 | 0.92 | 0.96 | 0.77 |
| Maxima during HiCo (MxCo) | 7-14 | .81 | .07 | 0.90 | 0.95 | 0.75 |
| Minima during LoCo (MnCo) | 6-17 | .41 | -.03 | 0.94 | 0.97 | 0.77 |
| Highest maxima (Mx) | 43 | .97 | .23** | 0.80** | 0.89** | 0.71** |
| Lowest minima (Mn) | 43 | .74 | .18* | 0.83* | 0.91* | 0.73* |

*Note:* Covariance maximizing eigenvector-based predictive modeling (CMEP; see Methods) was used in combination with a nested cross-validation scheme (see Methods, Fig. 1 and Fig. 2) to predict individual intelligence scores (latent *g*-factor derived from 12 cognitive scores) from static connectivity (all fMRI time frames; TFs), highest and lowest cofluctuation states (HiCo/LoCo; 43 TFs), maxima/minima *during* highest/lowest cofluctuation states (MxCo/MnCo; < 17 TFs), and the 43 highest maxima and lowest minima across the whole RSS time series (Mx, Mn; see Methods and Fig. 1). Reconstruction similarity values represent Pearson correlations between the static connectivity matrix (row 2) and the reconstructed connectivity matrix from the respective selection of time frames. Model performance metrics reflect the error between predicted and observed intelligence scores averaged across all four scans: Pearson correlation coefficient (*r*), mean squared error (MSE), root mean squared error (RMSE), and mean absolute error (MAE). Significance was determined by a non-parametric permutation test with 1,000 iterations for each scan and indicated as ** if $p < .001$, * if $p < .05$.

**Supplementary Tab. S3**

Prediction of intelligence from functional brain connectivity for the replication sample using the Yeo 114 nodes partition instead of Schaefer 100

| | TFs | Reconstruction similarity | *r* | MSE | RMSE | MAE |
|---|---|---|---|---|---|---|
| Static functional connectivity | 860 | n/a | .25** | 0.77** | 0.87** | 0.70** |
| High cofluctuations (HiCo) | 43 | .81 | .06 | 0.90 | 0.95 | 0.75 |
| Low cofluctuations (LoCo) | 43 | .52 | .06 | 0.88 | 0.94 | 0.75 |
| Maxima during HiCo (MxCo) | 7-14 | .81 | .05 | 0.90 | 0.95 | 0.75 |
| Minima during LoCo (MnCo) | 6-17 | .40 | -.01 | 0.91 | 0.96 | 0.76 |
| Highest Maxima (Mx) | 43 | .97 | .23** | 0.80** | 0.89** | 0.71** |
| Lowest Minima (Mn) | 43 | .74 | .17* | 0.83* | 0.91* | 0.73* |

*Note:* Covariance Maximizing Eigenvector-Based Predictive Modeling (CMEP; see Methods) was used in combination with a nested cross-validation scheme (see Methods, Fig. 1 and Fig 2) to predict individual intelligence scores (latent *g*-factor derived from 12 cognitive scores) from static connectivity (all fMRI time frames; TFs), highest and lowest cofluctuation states (HiCo/LoCo; 43 TFs), maxima/minima *during* highest/lowest cofluctuation states (MxCo/MnCo; < 17 TFs), and the 43 highest maxima and lowest minima across the whole RSS time series (Mx, Mn; see Methods and Fig. 1). Reconstruction similarity values represent Pearson correlations between the static connectivity matrix (row 2) and the reconstructed connectivity matrix from the respective selection of time frames. Model performance metrics reflect the error between predicted and observed intelligence scores averaged across all four scans: Pearson correlation coefficient (*r*), mean squared error (MSE), root mean squared error (RMSE), and mean absolute error (MAE). Significance was determined by a non-parametric permutation test with 1,000 iterations for each scan and indicated as ** if $p < .001$, * if $p < .05$ across all scans.
